# Prefrontal working memory activity partially mediates link between enriched neighborhood environments and episodic memory among 9-13-year-olds

**DOI:** 10.64898/2026.01.30.702968

**Authors:** Michael A. Rosario, Carlos Cardenas-Iniguez, Jennifer V. Chavez, Katherine Bottenhorn, Hedyeh Ahmadi, Megan M. Herting, Wesley K. Thompson

**Author notes:** Corresponding Author: **Dr. Michael A. Rosario**, University of Southern California, 1845 N. Soto St., Rm 225N, Los Angeles, CA, USA.

## Abstract

**Background:** Enriched environments support neurodevelopment. The pathways linking enrichment to cognitive processes such as episodic memory among youth remain unclear. This study examined whether brain function and structure in episodic memory-implicated neurocircuitry mediate the relationship between neighborhood enrichment and episodic memory performance.

**Methods:** We analyzed data from the Adolescent Brain Cognitive Development Study (n = 9,028) at two timepoints (baseline: 9-11-years-old and two-year follow-up: 11-13-years-old). Neighborhood enrichment was estimated at the child’s primary residential address proxied by the Child Opportunity Index 2.0 (COI). Episodic memory was assessed using the Picture Sequence Memory Test (PSMT). A multimodal neuroimaging approach examined task-based working memory-related functional activity, brain volume, and resting-state intrinsic activity in 26 bilateral brain regions, segmented using the Desikan-Killiany atlas, implicated in episodic memory. Following *FDR-*corrected linear mixed effects models, controlling for sociodemographic, neuroimaging factors, and site-related variability, two sets of mediation analyses were conducted per time point.

**Results:** Greater neighborhood enrichment (i.e., higher COI scores) was directly associated with better episodic memory, prefrontal cortex (PFC) task-based functional activity, and larger PFC and medial temporal lobe volume across timepoints. PFC task-based functional activity, but not brain volume or intrinsic activity, partially mediated these relationships. Specifically, PFC task-activity in the left and right caudal and rostral middle frontal gyri, and left pars opercularis, accounted for ∼2-7% of the mediated effect.

**Conclusion:** Our findings contribute to a rapidly growing body of literature linking environmental influences on neurocognitive outcomes during development. Given childhood and adolescence represent sensitive periods for neurodevelopment, interventions aimed at increasing neighborhood access to enriching experiences such as educational opportunities, cognitively stimulating activities, and social support may have lasting benefits for neurocognitive development.

## Introduction

Environmental enrichment has been widely regarded as a protective factor that buffers against the negative impacts of deprivation, fostering resilience and facilitating well-being(1). Conversely, environmental deprivation creates barriers to healthy development, detrimentally shaping long-term cognitive and neural outcomes(2). Beyond influencing cognitive ability, enrichment and deprivation may also shape functional and structural brain development over time. For example, experiences associated with enriched environments such as greater access to learning opportunities, cognitive stimulation, and social support are thought to shape the trajectory of structural and functional brain maturation, potentially supporting more efficient cortical networks in adulthood(3). In contrast, exposure to deprivation and adversity may accelerate brain development(4,5). These developmental differences underscore the importance of studying how environmental factors shape neurocognitive outcomes across childhood and adolescence.

The Child Opportunity Index 2.0 (COI), created by the Diversity Data Kids organization, provides a composite measure of census-tract level educational, socioeconomic, and health conditions that contribute to childhood development(6,7). Although the COI has been used to assess environmental influences on health and well-being(8–11), its impact on neurocognition, particularly during adolescent neurodevelopment, remains an important area of investigation. Unlike traditional socioeconomic status (SES) and socioeconomic position measures, the COI provides an encompassing assessment of environmental enrichment by capturing multiple dimensions of neighborhood-level resources including financial, health, environmental, and educational factors, allowing for a comprehensive analysis of how neighborhood enrichment affects cognitive and neural development.

A rich literature exists on the effects of environmental enrichment on brain function and structure using animal models(12,13). Rodent models of enrichment characterized by increased sensory, motor, and social stimulation have provided ample evidence of benefits to the brain and behavior including changes to brain plasticity(14), hippocampal neurogenesis(15), dendritic branching(16), and improvements to learning and memory(17–19). Comparatively in humans, the COI and other neighborhood level estimates, such as the Area Deprivation Index (ADI), have been used as proxies for neighborhood environmental enrichment or deprivation(9), respectively. Neuroimaging research has started to highlight how neighborhood-level enrichment, and its opposite, deprivation, may bolster or challenge the functional and structural neuroarchitecture involved in cognitive processes. Vargas and colleagues (2020) found that greater deprivation, measured using the ADI and linked to the child’s primary address, was associated with poorer cognition, comprising executive function, episodic memory, language, processing speed, working memory, and attention in late childhood(20). Moreover, in the same study greater deprivation was related to greater cortical thickness and surface area of the prefrontal cortex(20). In addition, higher ADI scores have been longitudinally linked to reduced episodic memory performance from ages 9-13(21), as well as cross-sectionally associated with smaller total hippocampal volume at 9-10-years-old(22) and smaller hippocampal subregion volumes(21). However, fewer studies have examined COI and its relation to childhood neurocognition. Recent work showed that lower COI (i.e., less neighborhood enrichment) was associated with greater prefrontal cortical thinning from ages 9-13(23). While these prior studies have established associations between deprivation or enrichment, cognition, and brain structure, the specific neural pathways that mediate the relationships between enrichment and neurocognition remain poorly understood. Understanding how enrichment shapes brain-behavior relationships during these sensitive periods is crucial for identifying the mechanisms that underlie cognitive development and for informing interventions aimed at optimizing learning and memory trajectories.

Overlapping cognitive systems essential for learning, memory, and identify formation undergo rapid maturation across childhood and adolescence(24). Episodic memory, the ability to recall specific life events, plays a fundamental role in both academic success and autobiographical identity, supporting learning and the integration of past experiences into personal narratives(25,26). Enriched environments, characterized by access to educational resources, social engagement, and physical activity, have been linked to better neurocognitive outcomes(27–29). However, while previous studies have demonstrated enrichment’s broad impact on cognition, little is known about how specific brain structures and functions mediate the relationship between enrichment and episodic memory. In particular, brain areas underlying learning and memory involve a complex, distributed network of regions within the prefrontal and parietal cortices and medial temporal lobes(30–33). The hippocampus, located within the medial temporal lobes, works in concert with regions of the parahippocampal gyrus in the formation and retrieval of episodic memories. Together, they contribute by integrating contextual information, which is crucial for the spatial and temporal aspects of episodic memory(33,34). Other regions, such as the precuneus of the parietal cortex, support the storage and retrieval of spatial and contextual information(35). Regions within the prefrontal cortex, including within the dorsolateral prefrontal cortex (dlPFC), are crucial for executive functions such as working memory, which is involved in regulating the organization of complex information during both encoding and retrieval of episodic memories(32,36). Recent work in adults has found dlPFC working memory-related functional activity to be associated with better episodic memory(37). Moreover, in childhood and adolescence episodic memory was longitudinally supported by a switch in recruitment from the hippocampus to the dlPFC(38), highlighting the importance of prefrontal functional activity on episodic memory. Together, these regions form a coordinated set of brain regions which support memory processes. Studies are thus needed to investigate how environmental enrichment impacts these pathways in order to clarify the neurobiological mechanisms that translate enriched experiences into cognitive benefits.

In this study, we examined the relationship between neighborhood environmental enrichment, as measured by COI, and episodic memory ability using the NIH Toolbox Picture Sequence Memory Test (PSMT). Additionally, we aimed to determine whether the functional and structural neural substrates of episodic memory mediate these relationships. Based on prior literature, we hypothesized that enriched neighborhood environments would positively influence brain function and structure, leading to better cognitive outcomes, specifically in learning and memory performance. Specifically, we expected greater task-related activation in memory-relevant brain regions (e.g., prefrontal cortex and medial temporal lobes) to mediate the relationship between neighborhood enrichment and episodic memory performance. This was based on prior literature identifying the relationship between working memory-related functional activity and episodic memory performance(37,38) and environmental factors with brain function and structure(4,21–23). We then extended these analyses to brain volume and resting state activity to determine whether the mediation effects were specific to task-evoked activity. By investigating these pathways, this study provides insight into the neural mechanisms linking enrichment to cognitive development, offering a more nuanced understanding of how environmental factors shape brain function.

## Methods

### Participants and ABCD Study Design

We used longitudinal data from the United States’ Adolescent Brain Cognitive Development^SM^ (ABCD) Study, an ongoing investigation into adolescent development. This analysis included a subset of data (**Supplemental Figure 1**) from the annual 5.1 data release (2023) (doi:10.15154/8873-zj65, NDA Study), including neuroimaging, cognitive, and linked external environmental data. The ABCD Study^®^ enrolled 11,880 children (between 2016 - 2018) aged 9 to 10 years from 21 sites across the United States and is following them longitudinally for ten years(39). The ABCD Study sample mirrors typical U.S. distributions for age, sex, and household size, but it underrepresents children of Asian, American Indian/Alaskan Native, and Native Hawaiian/Pacific Islander ancestry(40). Participants were excluded if they lacked English proficiency, had severe sensory, neurological, medical, or intellectual limitations, or were unable to complete an MRI scan. The ABCD study received approval from the Institutional Review Board (IRB) and Human Research Protections Programs (HRRP) at the University of California, San Diego. The IRB and HRRP at the University of California, San Diego oversee all experimental and consent procedures with local IRB board approval at the 21 ABCD sites. Written assent was provided by each participant, with written consent obtained from their legal guardians. For further details, see Garavan et al. (2018) and Volkow et al. (2018)(39,41). In this study, we analyzed a subset of the ABCD dataset, applying additional exclusion criteria based on MRI quality, incidental findings, and the availability of complete case data for our mediation analyses.

**Figure 1.**
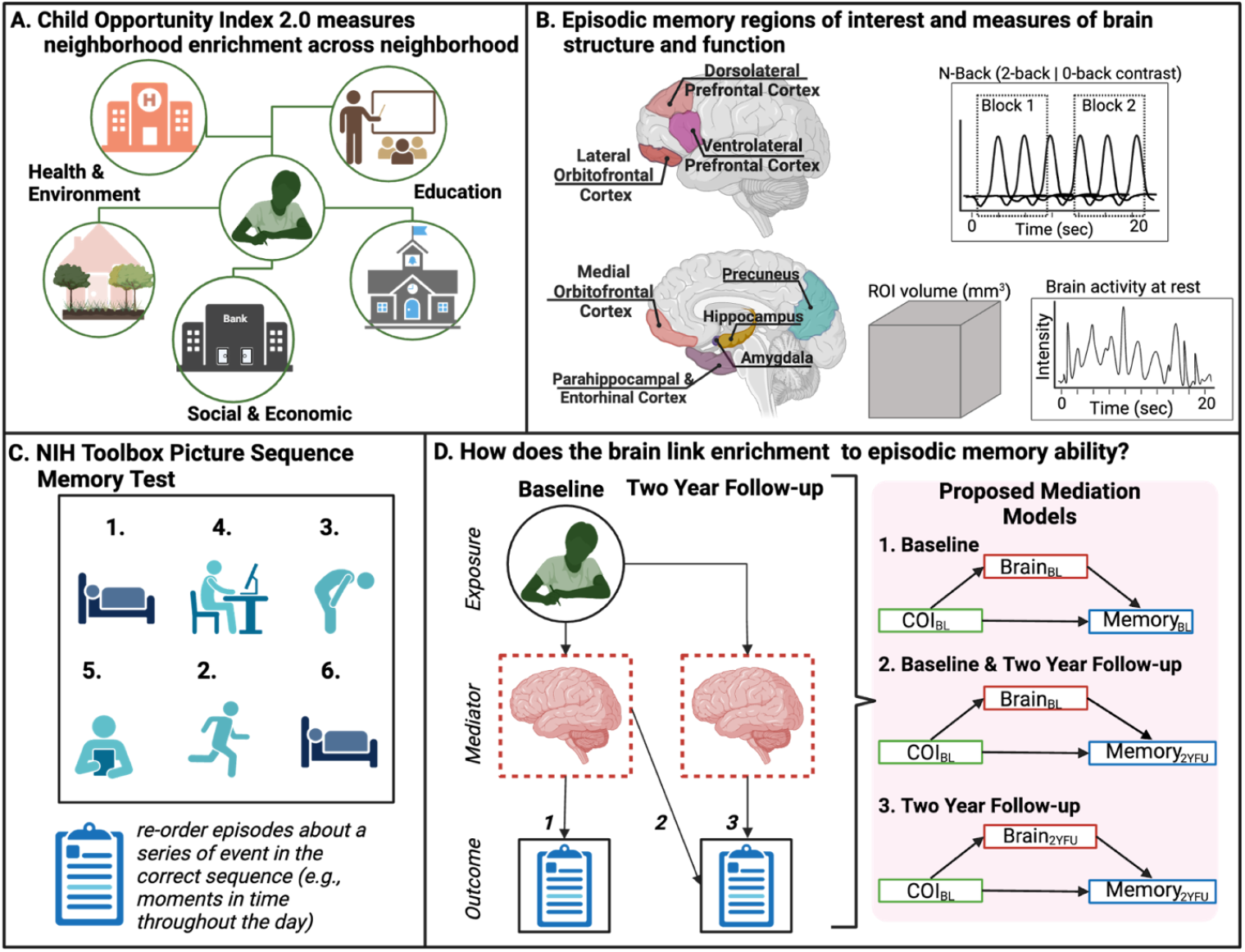
Overview of Childhood Opportunity Index 2.0, measures of brain regions important for episodic memory, picture sequence memory test ability, and mediation analysis breakdown. **A.** Details the Childhood Opportunity Index 2.0, comprising Health & Environment, Education, and Social & Economic domains. Ropresents resources available within the neighborhood which contribute to enriched child development. **B**, Provides a breakdown of brain regions important for episodic memory and their measures. **C**. Picture sequence memory test assesses an individual’s ability to remember and sequence pictures of events. **D**. Details analytical plan for mediation analyses, BL; baseline: COI; Childhood Opportunity Index 2.0; 2YFU: two year follow up.(Crealed in BioRender).

### Child Opportunity Index 2.0 (COI)

The COI was assigned to the primary residential addresses of each child at their baseline assessment(1,2)(Cardenas-Iniguez et al., 2024). It quantifies access to key childhood developmental resources across educational, health and environment, and social and economic domains (**Figure 1A**) with 29 key indicators of neighborhood resources measured at the census tract level(3,4). It reflects differences in neighborhood opportunities, with higher scores conceptualized as greater neighborhood enrichment. Some key indicators include: *Number of early childhood education centers within a 5-mile radius; Percent housing units that are vacant; Summer days with maximum temperature above 90F; Percent individuals living in households with incomes below 100% of the federal poverty threshold*(43). The tabulated COI z-score transformed composite score represents our primary exposure of interest.

### Regions of Interest

Regions of interest (ROIs) critical for episodic memory (**Figure 1B**), were extracted from the prefrontal cortex (*pars orbitalis, pars opercularis, pars triangularis, rostral caudal, caudal middle, and superior frontal gyri, medial and lateral orbitofrontal cortices*), parahippocampal gyrus (*parahippocampal and entorhinal cortices*), parietal cortex (*precuneus*), and subcortical regions (*hippocampus and amygdala*) using the Desikan-Killiany atlas(30,31,33,44). Motion correction and compliance procedures ensured high-quality data for downstream analysis. All ROI data were obtained from the ABCD tabulated dataset for brain volume, resting state functional magnetic resonance imaging (rs-fMRI), and task-based fMRI (t-fMRI)(5).

### Structural MRI: Acquisition, Processing, and Quality Control

Structural MRI protocols for the ABCD study have been described in detail previously(45,46). Briefly, harmonized protocols were implemented across sites using Siemens, Philips, or GE 3T MRI scanners. To minimize motion-related artifacts, participants underwent motion compliance training, and scans incorporated real-time, prospective motion correction. T1-weighted images (TE: 2–2.9 ms, TR: 6.31–2500 ms, T1: 1060 ms, flip angle: 8°, FOV: 256 x 256, resolution: 1 mm^3^ isotropic, 176 slices) were acquired using a magnetization-prepared rapid acquisition gradient echo (MPRAGE) sequence. Only images passing ABCD’s stringent quality-control parameters were included.

### Resting State fMRI (rs-fMRI) Temporal Variation: Acquisition, Processing, and Quality Control

Resting-state fMRI scans were collected using harmonized protocols on Siemens Prisma, Philips, or GE 750 3T MRI scanners(45). Imaging data acquisition included twenty cumulative minutes of resting-state data, split across four five-minute runs. Participants were instructed to keep their eyes open, focusing on a crosshair, to minimize variability in attentional state. To meet ABCD standards, scans included a minimum of 12.5 minutes of usable data with framewise displacement (FD) below 0.2 mm(47). Resting-state scans were acquired with an echo-planar imaging sequence in the axial plane (TR: 800 ms, TE: 30 ms, flip angle: 90°, voxel size: 2.4 mm^3^, 60 slices). Only images passing ABCD’s stringent quality-control parameters on low movement and no abnormal findings were included in analyses(46). Resting-state data were processed to extract time courses sampled onto cortical surfaces for each subject, using FreeSurfer-defined anatomical parcellations(44). Time courses for subcortical ROIs were calculated separately. Temporal variance, representing the magnitude of low-frequency fluctuations, its intrinsic activity(46), was computed for each ROI to serve as a measure of resting-state temporal variation. This metric provides insights into the non-task functional integrity of cortical and subcortical regions during rest.

### Task-based fMRI (t-fMRI): Acquisition, Processing, and Quality Control

Task-based fMRI data were acquired using standardized ABCD protocols and processed to control for motion artifacts and other confounds. Participants completed a working memory task (EN-Back), which utilized a block design alternating between 0-back and 2-back conditions. During the task, participants responded to repeated presentations of emotionally positive, negative, or neutral faces, as well as images of places(45,46,48). Nuisance regressors, such as motion estimates and derivatives, were included in the analysis, with time points showing framewise displacement (FD) over 0.9 mm censored. Beta coefficients and standard errors (SEM) were calculated for each ROI using AFNI’s 3dDeconvolve, incorporating stimulus timing and hemodynamic response functions. Runs with fewer than 50 degrees of freedom or extreme SEM outliers (>5% signal change) were excluded from group-level analyses to minimize the impact of motion variability, particularly among younger participants. After standard fMRI preprocessing, voxel time series were normalized, and time courses were sampled onto cortical surfaces using FreeSurfer. This approach projected gray matter voxels 1mm from the gray/white boundary. Average time courses were calculated for cortical and subcortical ROIs, as defined by the Desikan-Killiany atlas. Individual-level activation estimates were derived using a general linear model, providing ROI-based measures of working memory-related functional activity.

### Picture Sequence Memory Test (PSMT)

The NIH Toolbox PSMT was administered on a tablet and was used to measure nonverbal episodic memory(49). The test assesses an individual’s ability to remember and sequence pictures of events and shows good test-retest reliability(50). During the PSMT, participants are shown a series of pictures that tell a story, each presented one at a time in a specific order. Participants are then asked to recall the sequence by arranging the pictures in the correct order. Raw scores (derived from the cumulative number of adjacent pairs of pictures remembered correctly over three learning trials (e.g., 1-2-3-5-4-6; 2 points: 1-2 and 2-3) is the primary outcome of interest (**Figure 1C**).

### Statistical Analyses

All statistical analyses were conducted using R (v4.4.0). The primary objective was to determine whether measures of brain function and structure mediated the relationship between environmental enrichment, as indexed by the COI, and episodic memory performance during late childhood and early adolescence. The COI, assigned at baseline, was the primary exposure variable, while mediator variables included measures of brain function (working memory-related task-based activation and resting-state temporal variation) and structure (volume) from the hippocampus, amygdala, dlPFC, medial temporal lobes, and parietal cortex, assessed at both baseline and two-year follow-up. The outcome variables were raw scores from the PSMT, collected at both time points.

### Linear Mixed Effect Models

We first conducted a series of linear mixed-effects (LME) models to examine two key relationships. First, we assessed the association between enrichment (COI) and ROI-specific measures of brain function and structure. Next, we assessed the association between ROI-specific measures of brain function and structure and episodic memory performance. Since socio-demographic factors are correlated with COI and have been associated with neurodevelopmental outcomes, we included the following as covariates: race/ethnicity (White, Black, Hispanic, or Asian/Other), household income (>$100K, $50–100K, <$50K, or “Don’t Know/Refuse to Answer”), and highest parental education (Post-Graduate, Bachelor, Some College, High School Diploma/GED, or <High School Diploma), as well as sex at birth (male or female). Handedness (right, left, or mixed) and age (in months) were included as additional individual-level covariates. For MRI analyses, intracranial volume (ICV) was included as a covariate in structural MRI models, whereas mean framewise displacement for N-back task and resting state was included in task and resting-state fMRI models, respectively. We selected one child per family, and study site was included as a random effect to account for site-level differences in variables of interest. Following LME analyses, we applied false discovery rate (FDR) correction to account for multiple comparisons across 26 brain regions, ensuring that only significant relationships were carried forward to the mediation analyses.

### Mediation Analysis

For brain regions that showed significant relationships in the LME models, we next conducted multilevel causal mediation analyses using the *Mediation* package in R(51). This was done to examine whether brain function and structure mediated the relationship between enrichment and episodic memory performance. Mediation models were structured as follows: a) exposure variable: COI at baseline, b) mediator variables: brain measures significant after FDR-correction, and c) outcome variables: PSMT scores at baseline and two-year follow-up (**Figure 1D**). We ran three sets of mediation models to examine both cross-sectional and longitudinal relationships (two time points: baseline and two year follow-up) between enrichment, brain measures, and episodic memory performance. To ensure stability and robustness, each mediation analysis was conducted using 1,000 bootstrapped simulations to generate quasi-Bayesian confidence intervals for the average causal mediation effect (ACME), average direct effect (ADE), and proportion mediated. The same covariates from the LME models were included in all mediation analyses. This analytical approach allowed us to isolate the indirect effects of environmental enrichment on episodic memory performance through specific brain regions, while also assessing the direct influence of enrichment on memory outcomes.

## Results

The characteristics of the participant sample (n = 9,028; 48% female; 51% White; 9 - 13 years of age) are detailed in **Table 1**. A comparison between our analytical sample across analyses and the broader ABCD study cohort is provided in **Supplemental Table 1**. COI scores across the 21 study sites were relatively stable over analytical samples (**Supplemental Figures 2-4**), and PSMT scores improved from baseline to two year follow-up (**Table 1**).

|  | Baseline<br>(N=9028) | Baseline + Follow-Up<br>(N=8338) | Follow-Up<br>(N=5937) |
| --- | --- | --- | --- |
| **Age (months, Baseline)** |  |  |  |
| Mean (SD) | 120 (± 7.4) | 120 (± 7.4) | - |
| **Age (months, Follow-up)** |  |  |  |
| Mean (SD) | - | - | 140 (± 7.8) |
| **Sex** |  |  |  |
| Male | 4698 (52 %) | 4376 (52 %) | 3172 (53 %) |
| Female | 4330 (48 %) | 3962 (48 %) | 2765 (47 %) |
| **Race & Ethnicity** |  |  |  |
| Non-Hispanic White | 4605 (51 %) | 4366 (52 %) | 3218 (54 %) |
| Non-Hispanic Black | 1340 (15 %) | 1174 (14 %) | 793 (13 %) |
| Hispanic | 1928 (21 %) | 1734 (21 %) | 1189 (20 %) |
| Asian | 206 (2 %) | 188 (2 %) | 121 (2 %) |
| AIAN | 30 (0 %) | 25 (0 %) | 22 (0 %) |
| NHPI | 7 (0 %) | 6 (0 %) | 5 (0 %) |
| Other | 54 (1 %) | 49 (1 %) | 35 (1 %) |
| Multiracial Non-Black | 418 (5 %) | 392 (5 %) | 269 (5 %) |
| Multiracial Black | 392 (4 %) | 360 (4 %) | 259 (4 %) |
| DontKnow | 11 (0 %) | 9 (0 %) | 7 (0 %) |
| RefuseAnswer | 27 (0 %) | 25 (0 %) | 11 (0 %) |
| Missing | 10 (0.1%) | 10 (0.1%) | 8 (0.1%) |
| **Overall Income** |  |  |  |
| [<50K] | 2490 (28 %) | 2207 (26 %) | 1561 (26 %) |
| [≥50K and <100K] | 2334 (26 %) | 2199 (26 %) | 1634 (28 %) |
| [≥100K] | 3432 (38 %) | 3271 (39 %) | 2281 (38 %) |
| Don't know/Refuse to answer | 772 (9 %) | 661 (8 %) | 461 (8 %) |
| **Parental Highest Education** |  |  |  |
| < HS Diploma | 2239 (25 %) | 1986 (24 %) | 1389 (23 %) |
| HS Diploma/GED | 1111 (12 %) | 1001 (12 %) | 700 (12 %) |
| Some College | 1761 (20 %) | 1635 (20 %) | 1177 (20 %) |
| Bachelor Degree | 1997 (22 %) | 1898 (23 %) | 1351 (23 %) |
| Post Graduate Degree | 1813 (20 %) | 1726 (21 %) | 1250 (21 %) |
| Missing | 107 (1.2%) | 92 (1.1%) | 70 (1.2%) |
| **Picture Sequence Memory Test (Baseline)** |  |  |  |
| Mean (SD) | 100 (± 12) | 100 (± 12) | - |
| Missing | 120 (1.3%) | 105 (1.3%) | - |
| **Picture Sequence Memory Test (Follow-up)** |  |  |  |
| Mean (SD) | - | 110 (± 13) | 110 (± 12) |
| Missing | - | 444 (5.3%) | 214 (3.6%) |
| **COI Composite** |  |  |  |
| Mean (SD) | 0.011 (± 0.034) | 0.012 (± 0.034) | 0.012 (± 0.034) |
| Missing | 753 (8.3%) | 675 (8.1%) | 432 (7.3%) |

**Figure 2.**
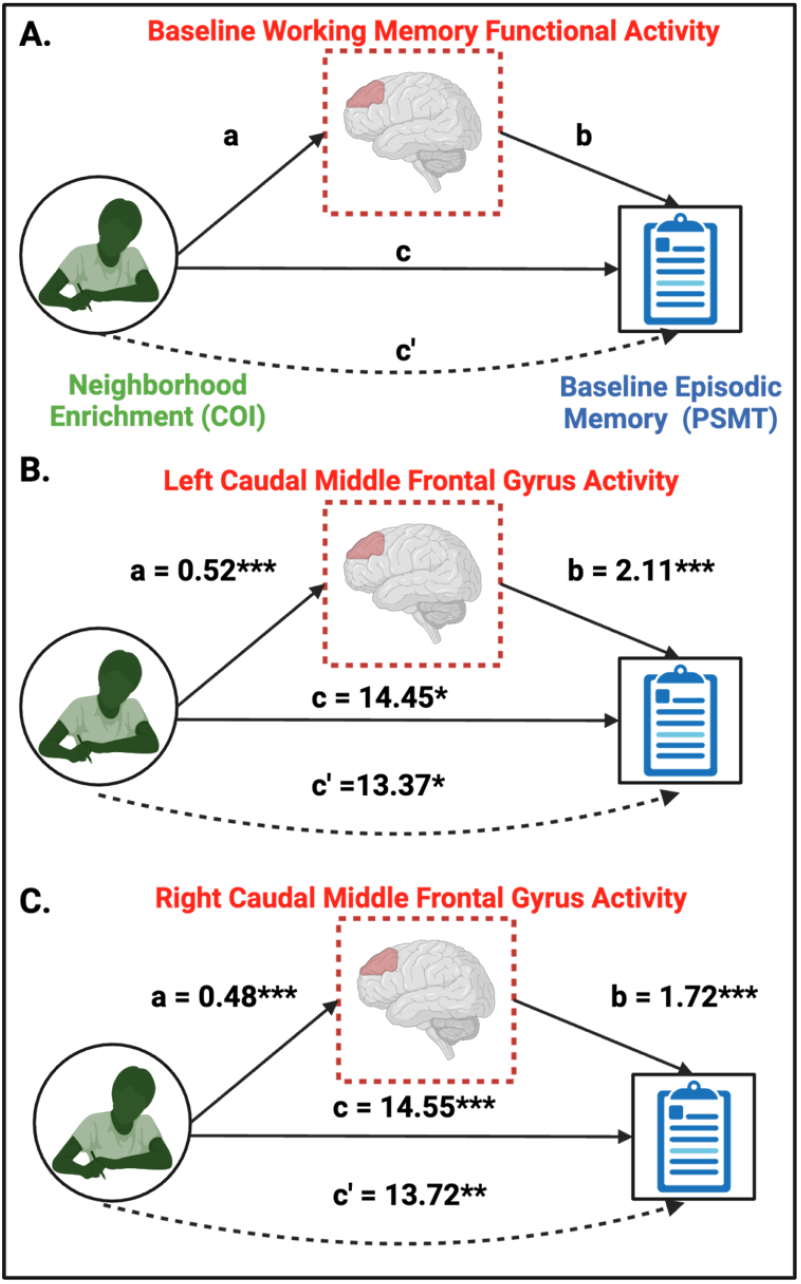
Prefrontal working-memory related activity links higher neighborhood enrichment with better episodic memory performance. A. Schematic overview of the mediation model, illustrating neighborhood enrichment (Child Opportunity Index; COI), baseline working memory-related functional activity, and baseline episodic memory performance (Picture Sequence Memory Test; PSMT). B. Left and C. right caudal middle frontal (CMF) gyrus working memory activity partially mediates higher neighborhood enrichment and better episodic memory performance. Dashed lines indicate indirect effects, while solid lines indicate direct relationships. Significance levels: ***p < 0.001, **p < 0.01, *p < 0.05.

**Figure 3.**
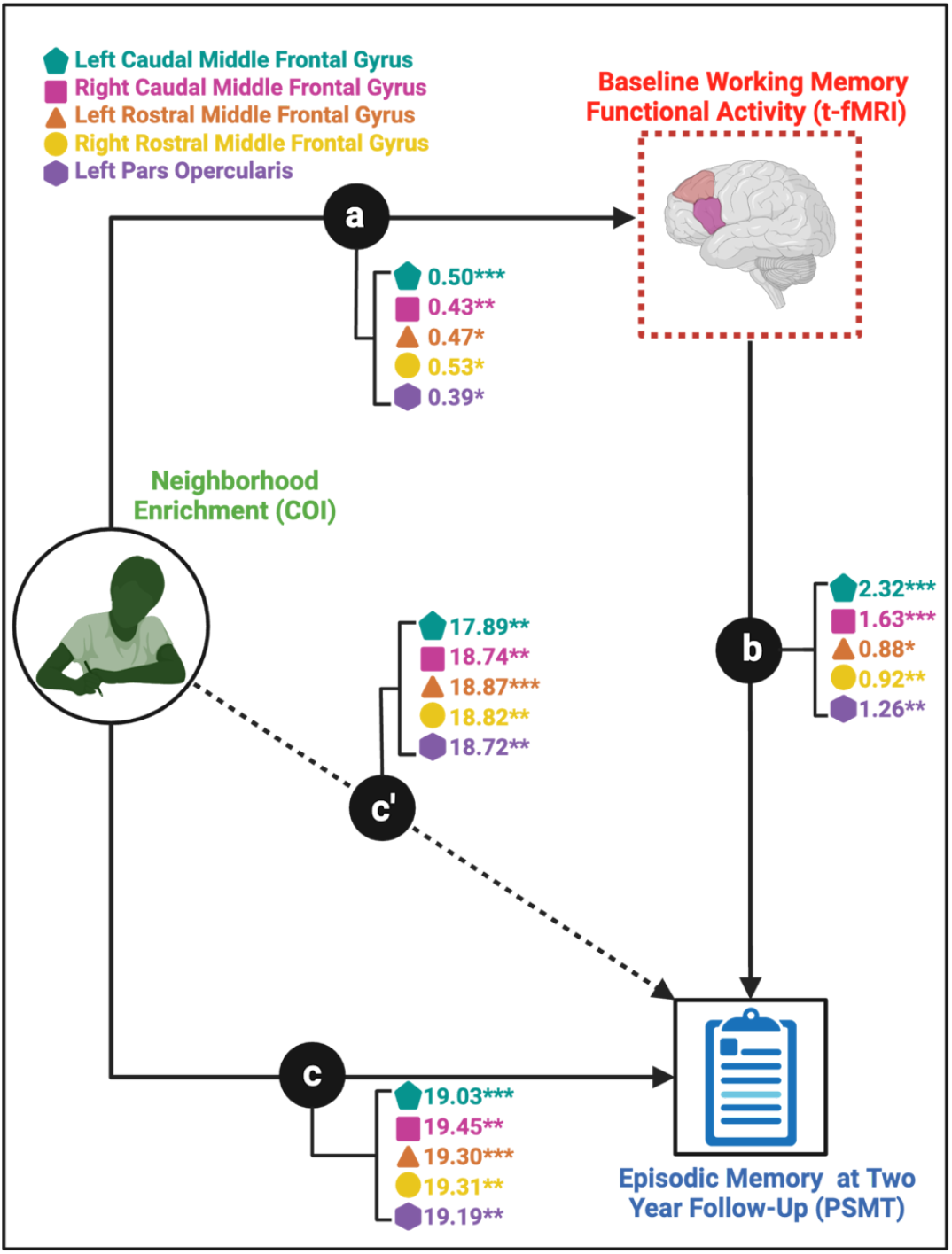
Prefrontal cortex working memory-related activity at baseline links enriched environments to episodic memory performance at follow-up. Path a represents the relationship between Child Opportunity Index (COI; enrichment) and baseline working memory-related functional activity in five prefrontal brain regions: left caudal middle frontal gyrus (CMF), right CMF, left rostral middle frontal gyrus (RMF), right RMF, and left pars opercularis. Path b represents the relationship between baseline working memory-related functional activity and episodic memory performance at the two-year follow-up (Picture Sequence Memory Test; PSMT). Path c’ represents the direct effect of COI on episodic memory performance, while path c represents the total effect (direct + indirect effects). Mediation analyses revealed that working memory-related activation in all five prefrontal regions significantly mediated the relationship between enrichment and episodic memory performance. The largest mediation effect was observed in the left CMF (5.9%), followed by right CMF (3.5%), left RMF (2.2%), right RMF (2.4%), and left pars opercularis (2.4%). These findings suggest that working memory-related activation in the prefrontal cortex partially explains the association between environmental enrichment and episodic memory outcomes during development. Significance levels: ***p < 0.001, **p < 0.01, *p < 0.05.

### Enrichment is related to episodic memory performance at baseline and at two year follow-up

In the linear mixed-effects model assessing the relationship between COI (enrichment) and PSMT scores (episodic memory performance) at baseline, higher COI scores were significantly associated with better PSMT scores (β = 14.83, 95%CI[3.93, 25.73], *p* = 0.008), controlling for sex, age, race and ethnicity, total household income, handedness, and motion, with study site (n= 21) as a random effect. In the subsequent model, additionally controlling for baseline PSMT scores, COI remained significantly associated with higher PSMT performance at the two-year follow-up (β = 18.74, 95%CI[7.76, 29.72], p < 0.001). In preparation for the final mediation model without baseline PSMT adjustment, the association between COI and episodic memory performance at follow-up was significant (β = 23.41, 95%CI[10.23, 36.59], *p* < 0.001).

### Enrichment is related to working memory functional activity and brain volume in regions of the parietal, mediotemporal, and prefrontal cortices

Linear mixed-effects models showed that environmental enrichment, proxied by COI, was significantly associated with baseline and two year follow-up working memory-related functional activation and cortical and subcortical volume in several brain regions implicated in episodic and working memory, after FDR correction (**Table 2**). In addition, measures of task-related activity and subcortical brain volume (at baseline) were associated with PSMT scores (**Table 3**). We then determined that of all linear mixed effect models, only left and right CMF, RMF, and left pars opercularis, met the criteria for mediation analyses for our first and second mediation models (i.e., PSMT at baseline and PSMT at two year follow-up) following FDR correction. In contrast, resting-state temporal variation was unrelated to both COI and PSMT performance at baseline and two year follow-up. For full baseline and two year follow-up LME model results, refer to **Supplemental Tables 4-21**).

**Table 2.** Linear mixed effect model results for all FDR-corrected significant relationships between neighborhood environmental enrichment (Child Opportunity Index 2.0) and task fMRI and volume ROIs across three sets of analyses (Baseline, Baseline + Follow-up, and Follow-up).

| ROI - Outcome | $\beta$ | Lower | Upper | *t* | *df* | *p* | *p adj* |
| --- | --- | --- | --- | --- | --- | --- | --- |
| *tfMRI - Baseline* |  |  |  |  |  |  |  |
| L Caudal Middle Frontal Gyrus | 0.48 | 0.22 | 0.74 | 3.59 | 3,106 | **<0.001** | **0.009** |
| R Caudal Middle Frontal Gyrus | 0.44 | 0.17 | 0.72 | 3.14 | 3,342 | **0.002** | **0.015** |
| L Rostral Middle Frontal Gyrus | 0.52 | 0.15 | 0.88 | 2.78 | 4,272 | **0.005** | **0.029** |
| R Rostral Middle Frontal Gyrus | 0.51 | 0.13 | 0.89 | 2.60 | 4,224 | **0.009** | **0.041** |
| L Precuneus | 0.43 | 0.17 | 0.69 | 3.23 | 2,825 | **0.001** | **0.015** |
| R Precuneus | 0.39 | 0.13 | 0.65 | 2.90 | 3,417 | **0.004** | **0.025** |
| *Volume - Baseline (mm<sup>3</sup>)* |  |  |  |  |  |  |  |
| L Amygdala | 230.48 | 75.20 | 385.80 | 2.91 | 8,241 | **0.004** | **0.023** |
| R Entorhinal Cortex | 422.19 | 104.77 | 739.60 | 2.61 | 5,408 | **0.009** | **0.04** |
| L Parahippocampus | 358.19 | 78.73 | 637.60 | 2.51 | 8,130 | **0.012** | **0.045** |
| L Rostral Middle Frontal Gyrus | 2,567.45 | 796.52 | 4338.40 | 2.84 | 8,265 | **0.005** | **0.023** |
| R Rostral Middle Frontal Gyrus | 3,405.28 | 1445.33 | 5365.20 | 3.41 | 8,272 | **0.001** | **0.012** |
| L Pars Orbitalis | 470.67 | 161.05 | 780.30 | 2.98 | 8,244 | **0.003** | **0.023** |
| R Pars Orbitalis | 622.75 | 254.43 | 991.10 | 3.31 | 8,200 | **0.001** | **0.012** |
| *tfMRI - Baseline and Follow-up* |  |  |  |  |  |  |  |
| R Caudal Middle Frontal Gyrus | 0.45 | 0.17 | 0.74 | 3.09 | 2,879 | **0.002** | **0.014** |
| L Rostral Middle Frontal Gyrus | 0.58 | 0.20 | 0.96 | 2.97 | 3,807 | **0.003** | **0.016** |
| R Rostral Middle Frontal Gyrus | 0.59 | 0.19 | 0.99 | 2.87 | 3,738 | **0.004** | **0.018** |
| L Pars Opercularis | 0.43 | 0.12 | 0.74 | 2.72 | 3,718 | **0.006** | **0.022** |
| R Pars Opercularis | 0.43 | 0.12 | 0.74 | 2.71 | 3,720 | **0.007** | **0.022** |
| L Precuneus | 0.48 | 0.20 | 0.75 | 3.40 | 2,627 | **0.001** | **0.009** |
| R Precuneus | 0.43 | 0.16 | 0.71 | 3.08 | 3,137 | **0.002** | **0.014** |
| *Volume - Baseline and Follow-up (mm<sup>3</sup>)* |  |  |  |  |  |  |  |
| L Amygdala | 207.75 | 45.18 | 370.31 | 2.50 | 7,545 | **0.012** | **0.047** |
| R Entorhinal Cortex | 424.86 | 91.06 | 758.66 | 2.49 | 5,015 | **0.013** | **0.047** |
| L Parahippocampus | 380.38 | 86.33 | 674.43 | 2.54 | 7,457 | **0.011** | **0.047** |
| L Rostral Middle Frontal Gyrus | 2,944.78 | 1,073.63 | 4,815.92 | 3.08 | 7,565 | **0.002** | **0.018** |
| R Rostral Middle Frontal Gyrus | 3,697.95 | 1,636.03 | 5,759.88 | 3.52 | 7,572 | **<0.001** | **0.006** |
| L Pars Orbitalis | 496.34 | 170.94 | 821.73 | 2.99 | 7,544 | **0.003** | **0.018** |
| R Pars Orbitalis | 701.63 | 312.16 | 1,091.10 | 3.53 | 7,498 | **<0.001** | **0.006** |
| *Volume - Follow-up (mm<sup>3</sup>)* |  |  |  |  |  |  |  |
| L Amygdala | 260.44 | 66.78 | 454.10 | 2.64 | 5,492 | **0.008** | **0.044** |
| R Amygdala | 290.48 | 114.07 | 466.88 | 3.23 | 5,175 | **0.001** | **0.011** |
| R Entorhinal Cortex | 654.71 | 280.32 | 1029.10 | 3.43 | 3,141 | **0.001** | **0.008** |
| L Parahippocampus | 441.00 | 96.11 | 785.90 | 2.51 | 5,381 | **0.012** | **0.05** |
| R Rostral Middle Frontal Gyrus | 4813.18 | 2419.09 | 7207.27 | 3.94 | 5,503 | **<0.001** | **0.002** |
| L Pars Orbitalis | 473.94 | 98.65 | 849.24 | 2.48 | 5,403 | **0.013** | **0.05** |
| R Pars Orbitalis | 654.81 | 201.80 | 1107.82 | 2.83 | 5,347 | **0.005** | **0.03** |

**Table 3.** Linear mixed effect model results for FDR-corrected significant relationship between episodic memory performance (PSMT scores) and tfMRI and volume ROIs across timepoints.

| ROI - Independent Variable | $\beta$ | Lower | Upper | *t* | *df* | *p* | *p adj* |
| --- | --- | --- | --- | --- | --- | --- | --- |
| *tfMRI - Baseline* |  |  |  |  |  |  |  |
| L Caudal Middle Frontal Gyrus | 2.14 | 1.22 | 3.05 | 4.56 | 7,523 | **<0.001** | **<0.001** |
| R Caudal Middle Frontal Gyrus | 1.65 | 0.78 | 2.52 | 3.72 | 7,524 | **<0.001** | **0.003** |
| *Volume (mm<sup>3</sup>) - Baseline* |  |  |  |  |  |  |  |
| L Hippocampus | 0.0016 | 0.0007 | 0.0024 | 3.71 | 8,511 | **<0.001** | **0.005** |
| *tfMRI - Baseline and Follow-up* |  |  |  |  |  |  |  |
| L Caudal Middle Frontal Gyrus | 2.36 | 1.40 | 3.32 | 4.83 | 6,619 | **<0.001** | **<0.001** |
| R Caudal Middle Frontal Gyrus | 1.42 | 0.52 | 2.33 | 3.09 | 6,620 | **0.002** | **0.018** |
| L Rostral Middle Frontal Gyrus | 1.01 | 0.32 | 1.71 | 2.86 | 6,617 | **0.004** | **0.025** |
| R Rostral Middle Frontal Gyrus | 0.95 | 0.29 | 1.60 | 2.82 | 6,614 | **0.005** | **0.025** |
| L Pars Opercularis | 1.43 | 0.57 | 2.29 | 3.26 | 6,614 | **0.001** | **0.015** |
| *tfMRI - Follow-up* |  |  |  |  |  |  |  |
| L Hippocampus | -2.03 | -3.22 | -0.84 | -3.34 | 5,155 | **0.001** | **0.004** |
| L Caudal Middle Frontal Gyrus | 3.27 | 2.18 | 4.36 | 5.89 | 5,155 | **<0.001** | **<0.001** |
| R Caudal Middle Frontal Gyrus | 3.43 | 2.34 | 4.51 | 6.19 | 5,155 | **<0.001** | **<0.001** |
| L Rostral Middle Frontal Gyrus | 1.78 | 0.87 | 2.68 | 3.85 | 5,156 | **<0.001** | **0.001** |
| R Rostral Middle Frontal Gyrus | 1.80 | 0.94 | 2.65 | 4.10 | 5,156 | **<0.001** | **<0.001** |
| L Pars Opercularis | 1.83 | 0.75 | 2.91 | 3.32 | 5,150 | **0.001** | **0.004** |

### Working memory functional activity in the prefrontal cortex mediates the relationship between enrichment and episodic memory performance at baseline and two year follow-up

Based on our linear mixed effects models, at baseline left and right CMF working memory-related functional activity met the criteria for mediation analysis (**Supplemental Table 2** for participant demographics). Our mediation analysis showed that both left and right CMF activity significantly mediated the relationship between COI and PSMT performance (**Figure 2**). For the left CMF, the ACME was statistically significant (β = 1.08, 95%CI[0.42, 1.95], *p* < 0.001) (**Figure 2A**). The ADE remained significant (β = 13.37, 95%CI[2.66, 24.75], *p* = 0.024). The total effect of COI on PSMT was significant (β = 14.45, 95%CI[3.68, 25.73], *p* = 0.022). The proportion of the total effect mediated by left CMF activity was 7.21% (*p* = 0.022). Similarly, for the right CMF, the ACME was also statistically significant (β = 1.72, 95%CI[0.80, 2.64], *p* < 0.001) (**Figure 2B**). The ADE remained significant (β = 13.72, 95%CI[2.80, 24.50], *p* = 0.014). The total effect of COI on PSMT was significant (β = 14.55, 95%CI[3.35, 25.75], *p* = 0.01). The proportion of the total effect mediated by right CMF activity was 4.79% (*p* < 0.001).

Following the same convention as our baseline analyses, we next tested whether working memory-related functional activity mediated the relationship between COI and PSMT scores (two year follow-up), while controlling for baseline PSMT scores see Supplemental Table 3 for participant demographics. Mediation analyses revealed that task-related functional activity in multiple prefrontal regions partially mediated the relationship between COI and PSMT performance (**Figure 3**). For the left CMF, the ACME was statistically significant (*β* = 1.14, 95%CI[0.44, 2.05], *p* < 0.001). The ADE was also significant (*β* = 17.89, 95%CI[6.81, 29.12], *p* = 0.002). The total effect of COI on memory was significant (*β* = 19.03, 95%CI[8.11, 30.31], *p* < 0.001), with 5.9% of this total effect mediated by left CMF activity. Similarly, the right CMF exhibited a significant mediation effect. The ACME was *β* = 0.71, 95%CI[0.18, 1.46], *p* = 0.006, with an ADE of *β* = 18.74, 95%CI[7.84, 29.25], *p* = 0.002 and a total effect of *β* = 19.45, 95%CI[8.57, 29.85], *p* = 0.002. The proportion of the total effect mediated was 3.5% (*p* = 0.008). For the left RMF, a small but significant mediation effect was observed. The ACME was *β* = 0.43, 95%CI[0.04, 1.04], *p* = 0.030, with an ADE of *β* = 18.86, 95%CI[8.31, 30.11], *p* = 0.002 and a total effect of *β* = 19.30, 95%CI[9.14, 30.75], *p* = 0.002. The proportion mediated was 2.2% (*p* = 0.032). For the right RMF, mediation was again present but modest in effect size. The ACME was *β* = 0.49, 95% CI[0.05, 1.12], *p* = 0.026, with an ADE of *β* = 18.82, 95%CI[7.42,29.54], *p* = 0.002 and a total effect of *β* = 19.31, 95%CI[7.63, 30.07], *p* = 0.002. The proportion mediated was 2.4% (*p* = 0.028). Finally, the left pars opercularis showed a significant mediation effect. The ACME was *β* = 0.48, 95%CI[0.04, 1.03], *p* = 0.022, with an ADE of *β* = 18.72, 95%CI[7.38, 29.76], *p* < 0.001, and a total effect of *β* = 19.19, 95%CI[7.81, 30.37], *p* < 0.001.The proportion mediated was 2.4% (*p* = 0.022).

## Discussion

This study examined the relationship between neighborhood environmental enrichment, brain function and structure, and episodic memory ability during late childhood and early adolescence. Using the COI as a proxy for environmental enrichment, we found that higher enrichment was positively associated with episodic memory performance, both at baseline and at the two-year follow-up. Enrichment was significantly associated with greater working memory related activity and greater cortical and subcortical volume across timepoints. Our mediation analyses revealed that baseline working memory-related activity in the PFC significantly mediated the relationship between enrichment and episodic memory performance measured at both timepoints. Across mediation models, task-based functional activity in the left and right CMF, left and right RMF, and left pars opercularis served as significant mediators between COI and episodic memory performance. In the first model, left and right CMF respectively accounted for 7.21% and 4.79% of the mediated effect linking enrichment and episodic memory performance at baseline. In our second model, the largest mediation effect was observed in the left CMF (5.9%), followed by the right CMF (3.5%), and smaller effects in the left RMF (2.2%), right RMF (2.4%), and left pars opercularis (2.4%) for episodic memory two years later. These results highlight the role of prefrontal cortex task engagement in linking enriched environments to episodic memory ability in late childhood and early adolescence, with CMF regions exhibiting the strongest effects. Notably, while mediation effects were present across all five regions, their magnitudes varied, suggesting region-specific contributions to the enrichment-memory relationship. These mediation effects were not observed when working memory-related function was measured at the two-year follow-up. These findings underscore the importance of enriched environments in shaping neurocognitive development and suggest that task-based functional activity may be particularly sensitive in explaining how environmental enrichment affects cognition.

### Environmental Enrichment and Episodic Memory

Our findings align with a growing body of research demonstrating that enriched environments are associated with enhanced cognitive outcomes in children(4,20–22,27,29). The present study extends these findings by providing evidence that enrichment is both associated with better episodic memory ability and that these cognitive benefits persist over time. Although we observed direct relationships between COI and both brain function and structure, only task-based functional activity mediated the relationship between COI and episodic memory performance. Neither brain volume nor resting-state connectivity played a significant mediating role. This suggests that functional brain activity during cognitive tasks may be a more direct or immediate mechanism over structure or resting-state(52) through which enrichment shapes memory ability. Since task-based activation reflects the engagement of memory-related neural circuits in real time, it may serve as a more sensitive indicator of enrichment’s effects on cognition compared to non-engaged measures of brain structure or intrinsic activity.

### Enrichment and Neurocognitive Development

Beyond task-based activity, our findings highlight the relationship between environmental enrichment and neurocognitive development. Previous research has shown that enrichment-related changes in brain function and structure play a crucial role in cognitive processes in childhood and adolescence(21,22,53). Additionally, animal models suggest that enriched environments promote neuroplasticity by increasing dendritic arborization, neurogenesis, and functional connectivity in key memory-related networks(12,15,16). In this study, COI was significantly associated with greater brain volume in memory-related regions. However, although episodic memory performance was significantly associated with left hippocampal volume (at baseline), there were no significant associations between brain volume and episodic memory performance. This suggests that enrichment may influence structural brain development without necessarily translating into cognitive benefits in the short term. Instead, brain volume alone may not be a sufficient predictor of cognitive function, instead relating to, for example, differences in childhood behavioral problems(9,10,54,55). In addition, enrichment-related volume earlier in life may support episodic memory during aging, influencing neurocognitive resilience and reserve(56–58). Future research should thus investigate the lifespan effects of early life enrichment. Here, we did not observe significant relationships between COI, episodic memory performance, and resting-state temporal activity across timepoints, suggesting that intrinsic brain activity at rest is not a significant mediator of enrichment effects on episodic memory. One possibility is that intrinsic functional activity may be more relevant in older adulthood in its link to episodic memory ability(59). Moreover, resting-state functional connectivity between brain regions, rather than the intrinsic activity within individual regions, may more directly reflect the neural mechanisms through which environmental enrichment influences cognitive outcomes(60).

### The Role of the PFC in the Enrichment-Episodic Memory Relationship

A key contribution of this study is the identification of working memory-related activity in the PFC as a mediator in the relationship between environmental enrichment and episodic memory performance. The PFC is a critical hub for executive function, supporting working memory(37,38). Previous work has shown that during late childhood and adolescence, episodic memory retrieval becomes increasingly dependent on prefrontal cortical engagement, reflecting a developmental shift from hippocampal to prefrontal-recruited memory systems(38). Our results suggest that enriched environments may enhance PFC function, which in turn supports episodic memory by facilitating working memory processes necessary for encoding and retrieval. Notably, although we found strong mediation effects for memory performance at baseline and at the two-year follow-up for baseline working memory task activity, there was no mediation effect when brain function was measured at follow-up. These findings were dependent on the lack of relationship between enrichment and working memory activity at follow-up. This suggests a potential proximal vs. distal temporal effect whereby task-based activity is potentially more temporally locked and a transient marker of neurocognitive plasticity in relation to enrichment. Over time, other genetic and developmental factors(55,61) may exert a greater influence on brain function and memory performance. Future research should investigate whether other factors beyond the neighborhood environment appreciably influence neurocognition in developing youth.

### Strengths & Limitations

One strength of this study is our use of the COI as a measure of neighborhood environmental enrichment. Much of the prior literature has examined environmental deprivation and SES as primary environmental influences on neurocognitive development. By using the COI, we were able to examine how neighborhood enrichment shapes brain function and episodic memory outcomes. Our findings also align with prior work on cognitive and brain reserve, which suggests that enriched experiences promote neural plasticity and functional adaptability, potentially fostering resilience against cognitive difficulties later in life(57,62). However, several limitations must be noted. The COI represents a relatively stable measure of neighborhood enrichment, but it does not capture day-to-day variability in a child’s lived experiences. Additionally, we did not examine how subdomains of the COI(3) relate to episodic memory. Future work should determine whether the separate COI subdomains differentially relate to episodic memory and if these relationships apply more broadly to other aspects of cognition.

## Acknowledgments

A special thank you to all participants and their families for their participation in the ABCD Study.

Data used in the preparation of this article were obtained from the Adolescent Brain Cognitive DevelopmentSM (ABCD) Study (https://abcdstudy.org), held in the NIMH Data Archive (NDA). This is a multisite, longitudinal study designed to recruit more than 10,000 children age 9-10 and follow them over 10 years into early adulthood. The ABCD Study® is supported by the National Institutes of Health and additional federal partners under award numbers U01DA041048, U01DA050989, U01DA051016, U01DA041022, U01DA051018, U01DA051037, U01DA050987, U01DA041174, U01DA041106, U01DA041117, U01DA041028, U01DA041134, U01DA050988, U01DA051039, U01DA041156, U01DA041025, U01DA041120, U01DA051038, U01DA041148,U01DA041093, U01DA041089, U24DA041123, U24DA041147. A full list of supporters is available at https://abcdstudy.org/federal-partners.html. A listing of participating sites and a complete listing of the study investigators can be found at https://abcdstudy.org/consortium_members/. ABCD consortium investigators designed and implemented the study and/or provided data but did not necessarily participate in the analysis or writing of this report. This manuscript reflects the views of the authors and may not reflect the opinions or views of the NIH or ABCD consortium investigators.

The ABCD data repository grows and changes over time. The ABCD data used in this report came from [doi:10.15154/8873-zj65]. DOIs can be found at [https://nda.nih.gov/study.html?id=2147].

## Author Contributions

Michael A. Rosario: Conceptualization, Formal Analysis, Writing - Original Draft and Editing, Visualization. Carlos Cardenas-Iniguez: Methodology, Writing – Review & Editing. Jennifer Chavez: Writing - Review & Editing. Katherine Bottenhorn: Writing - Review & Editing. Hedyeh Ahmadi: Statistical Supervision, Writing - Review & Editing. Megan M. Herting: Funding Acquisition, Methodology, Data Curation, Resources, Writing - Review & Editing. Wesley Thompson: Conceptualization, Methodology, Supervision, Project Administration, Writing - Review & Editing.

