## Supplemental Figures and Tables for "Prefrontal working memory activity partially mediates link between enriched neighborhood environments and episodic memory among 9-13-year-olds"

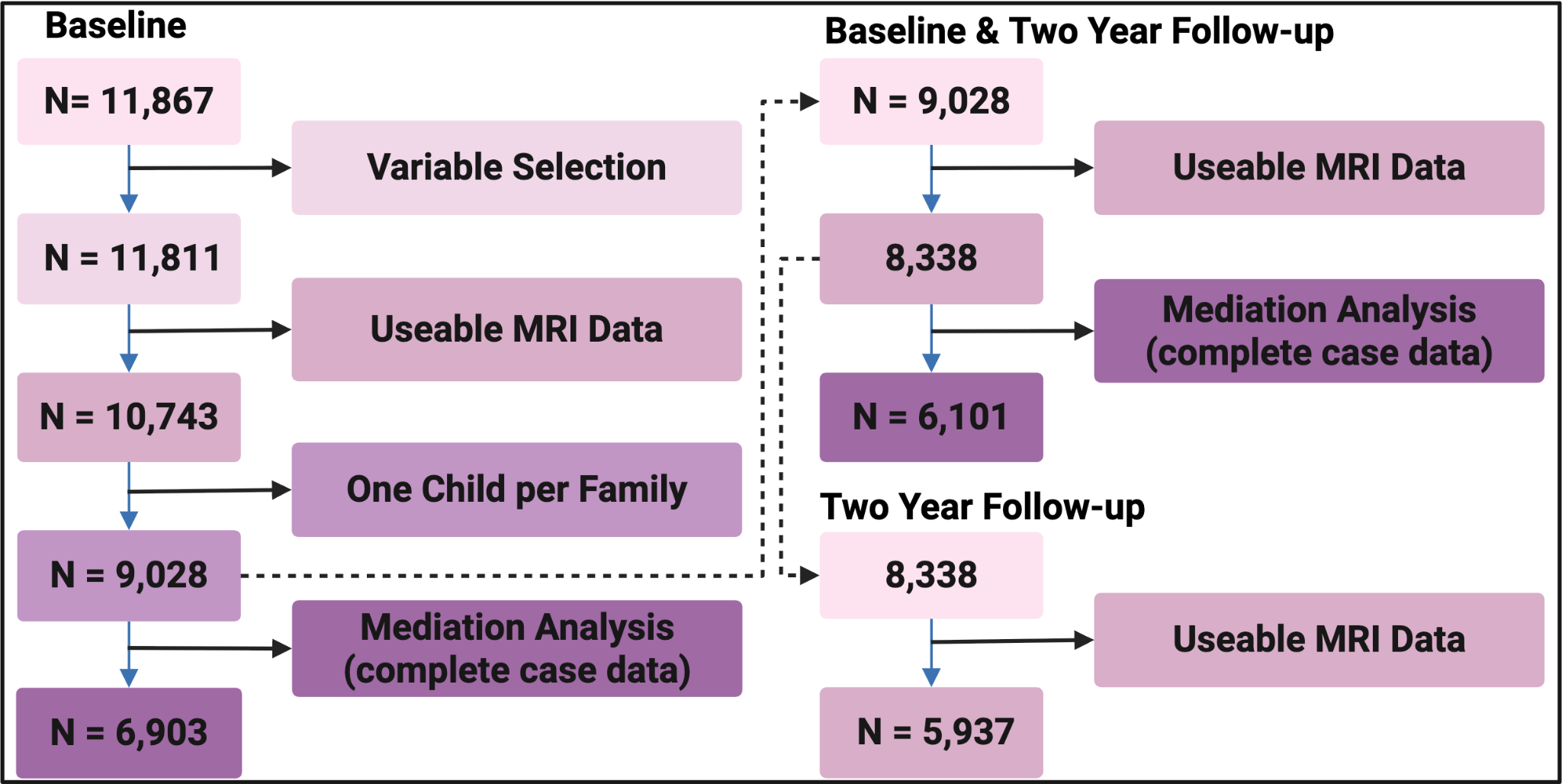

**S Fig 1. Participant exclusion flowchart for mediation analyses.** This diagram illustrates the sequential exclusion criteria applied to the study sample. Beginning with an initial sample of 11,867 participants at baseline, variable selection and data quality checks resulted in a subset with usable MRI data. Further exclusions ensured only one child per family was retained, reducing the sample to 9,028. Complete case data for mediation analysis at baseline resulted in a final sample of 6,903. For longitudinal analyses, baseline with two-year follow-up data were determined, yielding a final sample of 6,101 participants for mediation analyses. The flowchart also details exclusions for the two-year follow-up only, where 5,937 participants had usable MRI data.

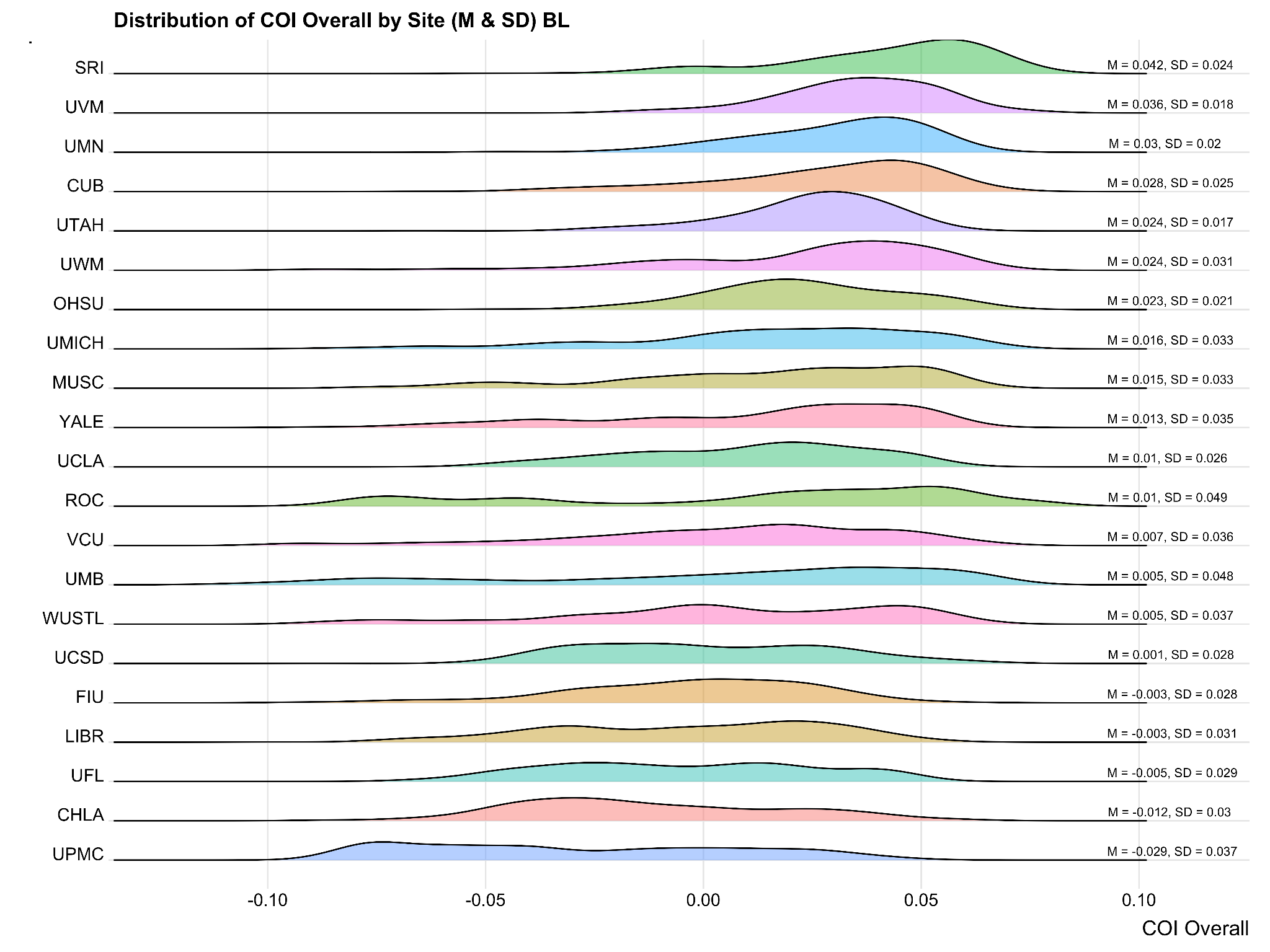

**S Figure 2.**  Distribution of Child Opportunity Index across ABCD Study Sites at Baseline Timepoint

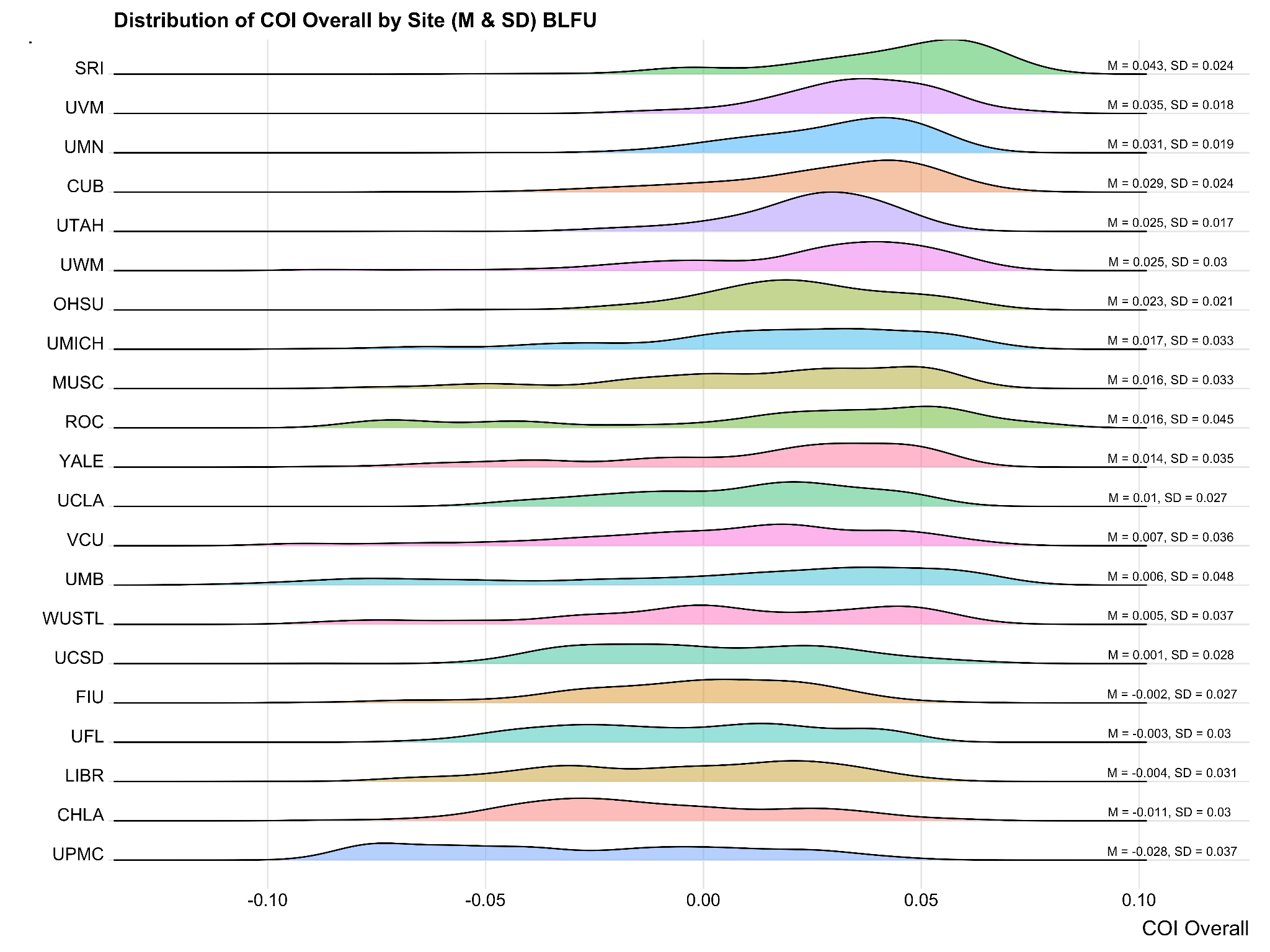

**S Figure 3.**  Distribution of Child Opportunity Index across ABCD Study Sites at Baseline and Follow-up Timepoints

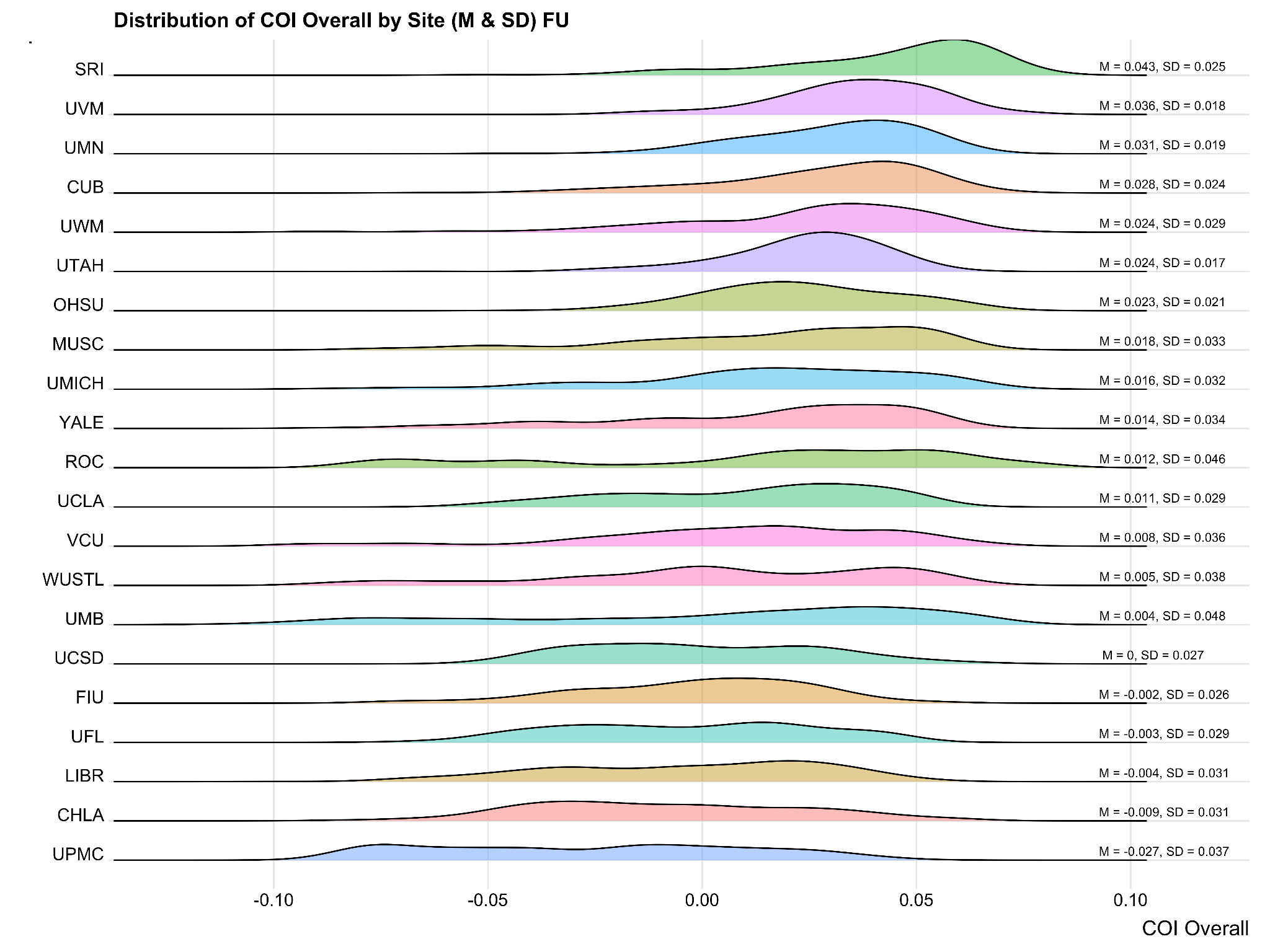

**S Figure 4.**  Distribution of Child Opportunity Index across ABCD Study Sites at Follow-up Timepoint

| **Supplemental Table 1** |  |  |  |  |
| --- | --- | --- | --- | --- |
|  | **Overall**  **(N=11867)** | **Baseline**  **(N=9028)** | **Baseline + Follow-Up**  **(N=8338)** | **Follow-Up**  **(N=5937)** |
| **Age (months, Baseline)** |  |  |  |  |
| Mean (SD) | 120 (± 7.5) | 120 (± 7.4) | 120 (± 7.4) | - |
| **Age (months, Follow-up)** | - | - | - |  |
| Mean (SD) | - | - | - | 140 (± 7.8) |
| **Sex** |  |  |  |  |
| Male | 6188 (52 %) | 4698 (52 %) | 4376 (52 %) | 3172 (53 %) |
| Female | 5676 (48 %) | 4330 (48 %) | 3962 (48 %) | 2765 (47 %) |
| Intersex-Male | 3 (0 %) | 0 (0 %) | 0 (0 %) | 0 (0 %) |
| **Race & Ethnicity** |  |  |  |  |
| Non-Hispanic White | 6167 (52 %) | 4605 (51 %) | 4366 (52 %) | 3218 (54 %) |
| Non-Hispanic Black | 1781 (15 %) | 1340 (15 %) | 1174 (14 %) | 793 (13 %) |
| Hispanic | 2410 (20 %) | 1928 (21 %) | 1734 (21 %) | 1189 (20 %) |
| Asian | 255 (2 %) | 206 (2 %) | 188 (2 %) | 121 (2 %) |
| AIAN | 39 (0 %) | 30 (0 %) | 25 (0 %) | 22 (0 %) |
| NHPI | 13 (0 %) | 7 (0 %) | 6 (0 %) | 5 (0 %) |
| Other | 66 (1 %) | 54 (1 %) | 49 (1 %) | 35 (1 %) |
| Multiracial Non-Black | 559 (5 %) | 418 (5 %) | 392 (5 %) | 269 (5 %) |
| Multiracial Black | 521 (4 %) | 392 (4 %) | 360 (4 %) | 259 (4 %) |
| DontKnow | 12 (0 %) | 11 (0 %) | 9 (0 %) | 7 (0 %) |
| RefuseAnswer | 30 (0 %) | 27 (0 %) | 25 (0 %) | 11 (0 %) |
| Missing | 14 (0.1%) | 10 (0.1%) | 10 (0.1%) | 8 (0.1%) |
| **Overall Income** |  |  |  |  |
| [<50K] | 3222 (27 %) | 2490 (28 %) | 2207 (26 %) | 1561 (26 %) |
| [≥50K and <100K] | 3067 (26 %) | 2334 (26 %) | 2199 (26 %) | 1634 (28 %) |
| [≥100K] | 4561 (38 %) | 3432 (38 %) | 3271 (39 %) | 2281 (38 %) |
| Don't know/Refuse to answer | 1015 (9 %) | 771 (9 %) | 661 (8 %) | 461 (8 %) |
| Missing | 2 (0.0%) | 1 (0.0%) | 0 (0%) | 0 (0%) |
| **Parental Highest Education** |  |  |  |  |
| < HS Diploma | 2893 (24 %) | 2239 (25 %) | 1986 (24 %) | 1389 (23 %) |
| HS Diploma/GED | 1449 (12 %) | 1111 (12 %) | 1001 (12 %) | 700 (12 %) |
| Some College | 2339 (20 %) | 1761 (20 %) | 1635 (20 %) | 1177 (20 %) |
| Bachelor Degree | 2655 (22 %) | 1997 (22 %) | 1898 (23 %) | 1351 (23 %) |
| Post Graduate Degree | 2381 (20 %) | 1813 (20 %) | 1726 (21 %) | 1250 (21 %) |
| Missing/Refused | 14 (0 %) | 0 (0 %) | 0 (0 %) | 0 (0 %) |
| Missing | 136 (1.1%) | 107 (1.2%) | 92 (1.1%) | 70 (1.2%) |
| **Picture Sequence Memory Test (Baseline)** | | |  |  |
| Mean (SD) | 100 (± 12) | 100 (± 12) | 100 (± 12) | - |
| Missing | 159 (1.3%) | 120 (1.3%) | 105 (1.3%) | - |
| **Picture Sequence Memory Test (Follow-up)** | | |  |  |
| Mean (SD) | - | - | 110 (± 13) | 110 (± 12) |
| Missing | - | - | 444 (5.3%) | 214 (3.6%) |
| **COI Composite** |  |  |  |  |
| Mean (SD) | 0.012 (± 0.034) | 0.011 (± 0.034) | 0.012 (± 0.034) | 0.012 (± 0.034) |
| Missing | 1092 (9.2%) | 753 (8.3%) | 675 (8.1%) | 432 (7.3%) |

**S Table 2 Descriptive Statistics for Mediation Analysis at Baseline**

|  | **Baseline**  **(N=6903)** |
| --- | --- |
| **Age (months, Baseline)** |  |
| Mean (SD) | 120 (± 7.5) |
| **Sex** |  |
| Male | 3530 (51 %) |
| Female | 3373 (49 %) |
| **Race & Ethnicity** |  |
| White | 3674 (53 %) |
| Black | 886 (13 %) |
| Hispanic | 1455 (21 %) |
| Asian/Other | 888 (13 %) |
| **Overall Income** |  |
| [<50K] | 1759 (25 %) |
| [≥50K and <100K] | 1828 (26 %) |
| [≥100K] | 2752 (40 %) |
| Don't know/Refuse to answer | 564 (8 %) |
| **Parental Highest Education** |  |
| < HS Diploma | 1613 (23 %) |
| HS Diploma/GED | 791 (11 %) |
| Some College | 1359 (20 %) |
| Bachelor Degree | 1617 (23 %) |
| Post Graduate Degree | 1448 (21 %) |
| Missing/Refused | 0 (0 %) |
| Missing | 75 (1.1%) |
| **Picture Sequence Memory Test** |  |
| Mean (SD) | 100 (± 12) |
| **COI Composite** |  |
| Mean (SD) | 0.013 (± 0.033) |

**S Table 3 Descriptive Statistics for Mediation Analysis at Baseline and**

**Two Year Follow-Up**

|  | **Baseline + Follow-Up**  **(N=6101)** |
| --- | --- |
| **Age (months, Baseline)** |  |
| Mean (SD) | 120 (± 7.4) |
| **Sex** |  |
| Male | 3139 (51 %) |
| Female | 2962 (49 %) |
| **Race & Ethnicity** |  |
| White | 3317 (54 %) |
| Black | 730 (12 %) |
| Hispanic | 1266 (21 %) |
| Asian/Other | 788 (13 %) |
| **Overall Income** |  |
| [<50K] | 1504 (25 %) |
| [≥50K and <100K] | 1637 (27 %) |
| [≥100K] | 2495 (41 %) |
| Don't know/Refuse to answer | 465 (8 %) |
| **Parental Highest Education** |  |
| < HS Diploma | 1383 (23 %) |
| HS Diploma/GED | 674 (11 %) |
| Some College | 1200 (20 %) |
| Bachelor Degree | 1477 (24 %) |
| Post Graduate Degree | 1303 (21 %) |
| Missing/Refused | 0 (0 %) |
| Missing | 64 (1.0%) |
| **Picture Sequence Memory Test (Follow)** | |
| Mean (SD) | 110 (± 12) |
| **Picture Sequence Memory Test (Baseline)** | |
| Mean (SD) | 100 (± 12) |
| **COI Composite** |  |
| Mean (SD) | 0.014 (± 0.033) |

### **S Table 4.** Linear mixed effect model results for relationship between enrichment and tfMRI ROI at Baseline

| ROI - Outcome | 𝛽 | Lower | Upper | t | df | p | p adj |
| --- | --- | --- | --- | --- | --- | --- | --- |
| L Hippocampus | 0.06 | -0.23 | 0.34 | 0.39 | 2,370 | 0.698 | 0.841 |
| R Hippocampus | 0.04 | -0.25 | 0.33 | 0.24 | 3,086 | 0.808 | 0.841 |
| L Amygdala | 0.04 | -0.36 | 0.45 | 0.20 | 2,309 | 0.841 | 0.841 |
| R Amygdala | -0.30 | -0.71 | 0.12 | -1.41 | 2,786 | 0.159 | 0.318 |
| L Entorhinal Cortex | 0.16 | -0.87 | 1.19 | 0.30 | 2,386 | 0.762 | 0.841 |
| R Entorhinal Cortex | -0.21 | -1.57 | 1.16 | -0.30 | 2,109 | 0.767 | 0.841 |
| L Parahippocampus | -0.17 | -0.54 | 0.20 | -0.92 | 3,059 | 0.359 | 0.549 |
| R Parahippocampus | -0.26 | -0.66 | 0.13 | -1.31 | 2,897 | 0.191 | 0.353 |
| L Lateral Orbitofrontal Cortex | 0.14 | -0.40 | 0.68 | 0.52 | 2,872 | 0.601 | 0.822 |
| R Lateral Orbitofrontal Cortex | 0.25 | -0.34 | 0.84 | 0.83 | 2,674 | 0.408 | 0.589 |
| L Medial Orbitofrontal Cortex | 0.08 | -0.62 | 0.78 | 0.23 | 2,529 | 0.820 | 0.841 |
| R Medial Orbitofrontal Cortex | 0.07 | -0.61 | 0.76 | 0.21 | 6,980 | 0.837 | 0.841 |
| L Caudal Middle Frontal Gyrus | 0.48 | 0.22 | 0.74 | 3.59 | 3,106 | **<0.001** | **0.009** |
| R Caudal Middle Frontal Gyrus | 0.44 | 0.17 | 0.72 | 3.14 | 3,342 | **0.002** | **0.015** |
| L Rostral Middle Frontal Gyrus | 0.52 | 0.15 | 0.88 | 2.78 | 4,272 | **0.005** | **0.029** |
| R Rostral Middle Frontal Gyrus | 0.51 | 0.13 | 0.89 | 2.60 | 4,224 | **0.009** | **0.041** |
| L Pars Opercularis | 0.33 | 0.04 | 0.63 | 2.19 | 4,153 | **0.028** | 0.093 |
| R Pars Opercularis | 0.37 | 0.07 | 0.66 | 2.41 | 4,159 | **0.016** | 0.060 |
| L Pars Triangularis | 0.37 | 0.02 | 0.71 | 2.07 | 2,788 | **0.039** | 0.112 |
| R Pars Triangularis | 0.26 | -0.10 | 0.63 | 1.44 | 3,199 | 0.150 | 0.318 |
| L Pars Orbitalis | 0.43 | -0.38 | 1.24 | 1.04 | 2,992 | 0.296 | 0.482 |
| R Pars Orbitalis | 0.53 | -0.29 | 1.34 | 1.27 | 3,007 | 0.204 | 0.353 |
| L Superior Frontal Gyrus | 0.24 | 0.00 | 0.48 | 1.92 | 2,645 | 0.055 | 0.143 |
| R Superior Frontal Gyrus | 0.24 | -0.01 | 0.48 | 1.85 | 3,038 | 0.065 | 0.153 |
| L Precuneus | 0.43 | 0.17 | 0.69 | 3.23 | 2,825 | **0.001** | **0.015** |
| R Precuneus | 0.39 | 0.13 | 0.65 | 2.90 | 3,417 | **0.004** | **0.025** |

### **S Table 5.** Linear mixed effect model results for relationship between PSMT scores and tfMRI ROI at Baseline

| ROI - Predictor | 𝛽 | Lower | Upper | t | df | p | p adj |
| --- | --- | --- | --- | --- | --- | --- | --- |
| L Hippocampus | -0.45 | -1.31 | 0.41 | -1.03 | 7,520 | 0.302 | 0.466 |
| R Hippocampus | -0.59 | -1.42 | 0.23 | -1.41 | 7,523 | 0.159 | 0.464 |
| L Amygdala | -0.13 | -0.72 | 0.47 | -0.42 | 7,519 | 0.673 | 0.833 |
| R Amygdala | -0.33 | -0.91 | 0.25 | -1.11 | 7,521 | 0.268 | 0.464 |
| L Entorhinal Cortex | -0.15 | -0.39 | 0.10 | -1.19 | 7,524 | 0.235 | 0.464 |
| R Entorhinal Cortex | -0.02 | -0.20 | 0.16 | -0.25 | 7,520 | 0.800 | 0.905 |
| L Parahippocampus | -0.38 | -1.04 | 0.29 | -1.12 | 7,522 | 0.264 | 0.464 |
| R Parahippocampus | -0.47 | -1.08 | 0.14 | -1.52 | 7,522 | 0.130 | 0.464 |
| L Lateral Orbitofrontal Cortex | 0.24 | -0.21 | 0.68 | 1.03 | 7,519 | 0.304 | 0.466 |
| R Lateral Orbitofrontal Cortex | -0.06 | -0.48 | 0.35 | -0.30 | 7,520 | 0.765 | 0.904 |
| L Medial Orbitofrontal Cortex | -0.23 | -0.57 | 0.12 | -1.29 | 7,518 | 0.198 | 0.464 |
| R Medial Orbitofrontal Cortex | -0.27 | -0.62 | 0.08 | -1.52 | 7,517 | 0.129 | 0.464 |
| L Caudal Middle Frontal Gyrus | 2.14 | 1.22 | 3.05 | 4.56 | 7,523 | **<0.001** | **<0.001** |
| R Caudal Middle Frontal Gyrus | 1.65 | 0.78 | 2.52 | 3.72 | 7,524 | **<0.001** | **0.003** |
| L Rostral Middle Frontal Gyrus | 0.73 | 0.07 | 1.39 | 2.18 | 7,525 | **0.029** | 0.187 |
| R Rostral Middle Frontal Gyrus | 0.67 | 0.04 | 1.29 | 2.10 | 7,525 | **0.036** | 0.187 |
| L Pars Opercularis | 1.15 | 0.33 | 1.98 | 2.74 | 7,526 | **0.006** | 0.053 |
| R Pars Opercularis | 0.54 | -0.27 | 1.36 | 1.31 | 7,525 | 0.191 | 0.464 |
| L Pars Triangularis | 0.02 | -0.67 | 0.71 | 0.07 | 7,522 | 0.947 | 0.976 |
| R Pars Triangularis | -0.44 | -1.10 | 0.22 | -1.31 | 7,521 | 0.191 | 0.464 |
| L Pars Orbitalis | -0.18 | -0.48 | 0.13 | -1.14 | 7,521 | 0.253 | 0.464 |
| R Pars Orbitalis | -0.12 | -0.41 | 0.18 | -0.77 | 7,521 | 0.443 | 0.641 |
| L Superior Frontal Gyrus | -0.04 | -1.02 | 0.93 | -0.08 | 7,521 | 0.933 | 0.976 |
| R Superior Frontal Gyrus | 0.25 | -0.70 | 1.20 | 0.51 | 7,522 | 0.608 | 0.791 |
| L Precuneus | 0.01 | -0.92 | 0.95 | 0.03 | 7,523 | 0.976 | 0.976 |
| R Precuneus | 0.32 | -0.62 | 1.25 | 0.67 | 7,525 | 0.505 | 0.690 |

### **S Table 6.** Linear mixed effect model results for relationship between enrichment and ROI volume at Baseline

| ROI - Outcome | 𝛽 | Lower | Upper | t | df | p | p adj |
| --- | --- | --- | --- | --- | --- | --- | --- |
| L Hippocampus | 155.33 | -99.03 | 409.7 | 1.20 | 8,073 | 0.231 | 0.384 |
| R Hippocampus | 314.63 | 42.23 | 587.0 | 2.26 | 7,670 | **0.024** | 0.077 |
| L Amygdala | 230.48 | 75.20 | 385.8 | 2.91 | 8,241 | **0.004** | **0.023** |
| R Amygdala | 160.36 | 17.26 | 303.4 | 2.20 | 7,744 | **0.028** | 0.081 |
| L Entorhinal Cortex | 251.12 | -75.74 | 578.0 | 1.51 | 8,090 | 0.132 | 0.245 |
| R Entorhinal Cortex | 422.19 | 104.77 | 739.6 | 2.61 | 5,408 | **0.009** | **0.040** |
| L Parahippocampus | 358.19 | 78.73 | 637.6 | 2.51 | 8,130 | **0.012** | **0.045** |
| R Parahippocampus | 238.41 | 8.24 | 468.6 | 2.03 | 7,159 | **0.042** | 0.100 |
| L Lateral Orbitofrontal Cortex | 674.49 | 35.99 | 1,313.0 | 2.07 | 8,272 | **0.038** | 0.100 |
| R Lateral Orbitofrontal Cortex | 599.13 | -55.78 | 1,254.0 | 1.79 | 8,264 | 0.073 | 0.158 |
| L Medial Orbitofrontal Cortex | -44.93 | -548.25 | 458.4 | -0.17 | 8,261 | 0.861 | 0.896 |
| R Medial Orbitofrontal Cortex | -108.48 | -623.55 | 406.6 | -0.41 | 8,271 | 0.680 | 0.842 |
| L Caudal Middle Frontal Gyrus | 653.69 | -435.51 | 1,742.9 | 1.18 | 8,272 | 0.240 | 0.384 |
| R Caudal Middle Frontal Gyrus | -398.65 | -1,501.69 | 704.4 | -0.71 | 8,271 | 0.479 | 0.655 |
| L Rostral Middle Frontal Gyrus | 2,567.45 | 796.52 | 4,338.4 | 2.84 | 8,265 | **0.005** | **0.023** |
| R Rostral Middle Frontal Gyrus | 3,405.28 | 1,445.33 | 5,365.2 | 3.41 | 8,272 | **0.001** | **0.012** |
| L Pars Opercularis | -91.26 | -893.39 | 710.9 | -0.22 | 8,137 | 0.824 | 0.892 |
| R Pars Opercularis | 92.98 | -554.70 | 740.6 | 0.28 | 8,169 | 0.778 | 0.884 |
| L Pars Triangularis | -157.41 | -805.02 | 490.2 | -0.48 | 8,041 | 0.634 | 0.824 |
| R Pars Triangularis | -342.32 | -1,095.63 | 411.0 | -0.89 | 8,159 | 0.373 | 0.539 |
| L Pars Orbitalis | 470.67 | 161.05 | 780.3 | 2.98 | 8,244 | **0.003** | **0.023** |
| R Pars Orbitalis | 622.75 | 254.43 | 991.1 | 3.31 | 8,200 | **0.001** | **0.012** |
| L Superior Frontal Gyrus | -28.22 | -2,191.21 | 2,134.8 | -0.03 | 8,272 | 0.980 | 0.980 |
| R Superior Frontal Gyrus | 308.62 | -1,872.79 | 2,490.0 | 0.28 | 8,270 | 0.782 | 0.884 |
| L Precuneus | 614.89 | -435.19 | 1,665.0 | 1.15 | 8,263 | 0.251 | 0.384 |
| R Precuneus | 862.23 | -204.47 | 1,928.9 | 1.58 | 8,273 | 0.113 | 0.226 |

### **S Table 7.** Linear mixed effect model results for relationship between PSMT scores and ROI volume at Baseline

| ROI - Predictor | 𝛽 | Lower | Upper | t | df | p | p adj |
| --- | --- | --- | --- | --- | --- | --- | --- |
| L Hippocampus | 0.0016 | 0.0007 | 0.0024 | 3.71 | 8,511 | **<0.001** | **0.005** |
| R Hippocampus | 0.0010 | 0.0002 | 0.0018 | 2.57 | 8,798 | **0.010** | 0.089 |
| L Amygdala | 0.0003 | -0.0011 | 0.0017 | 0.43 | 7,817 | 0.666 | 0.862 |
| R Amygdala | -0.0003 | -0.0018 | 0.0011 | -0.44 | 8,793 | 0.662 | 0.862 |
| L Entorhinal Cortex | 0.0007 | 0.0000 | 0.0013 | 2.10 | 8,534 | **0.036** | 0.187 |
| R Entorhinal Cortex | 0.0005 | -0.0002 | 0.0011 | 1.40 | 8,906 | 0.162 | 0.383 |
| L Parahippocampus | 0.0007 | 0.0000 | 0.0014 | 1.83 | 8,424 | 0.067 | 0.219 |
| R Parahippocampus | 0.0001 | -0.0008 | 0.0010 | 0.11 | 8,874 | 0.911 | 0.925 |
| L Lateral Orbitofrontal Cortex | 0.0003 | 0.0000 | 0.0007 | 2.11 | 5,255 | **0.035** | 0.187 |
| R Lateral Orbitofrontal Cortex | 0.0004 | 0.0001 | 0.0007 | 2.66 | 955 | **0.008** | 0.089 |
| L Medial Orbitofrontal Cortex | 0.0000 | -0.0004 | 0.0003 | -0.24 | 741 | 0.807 | 0.887 |
| R Medial Orbitofrontal Cortex | 0.0002 | -0.0002 | 0.0006 | 0.83 | 3,275 | 0.409 | 0.755 |
| L Caudal Middle Frontal Gyrus | 0.0000 | -0.0002 | 0.0002 | 0.23 | 5,707 | 0.819 | 0.887 |
| R Caudal Middle Frontal Gyrus | 0.0001 | -0.0001 | 0.0003 | 0.89 | 5,836 | 0.373 | 0.746 |
| L Rostral Middle Frontal Gyrus | 0.0000 | -0.0001 | 0.0001 | 0.29 | 6,644 | 0.774 | 0.887 |
| R Rostral Middle Frontal Gyrus | 0.0000 | -0.0001 | 0.0001 | 0.78 | 5,403 | 0.436 | 0.755 |
| L Pars Opercularis | -0.0001 | -0.0003 | 0.0002 | -0.43 | 8,505 | 0.665 | 0.862 |
| R Pars Opercularis | -0.0003 | -0.0006 | 0.0000 | -1.77 | 8,413 | 0.076 | 0.220 |
| L Pars Triangularis | -0.0001 | -0.0004 | 0.0002 | -0.44 | 8,627 | 0.658 | 0.862 |
| R Pars Triangularis | 0.0001 | -0.0002 | 0.0003 | 0.39 | 8,432 | 0.696 | 0.862 |
| L Pars Orbitalis | 0.0007 | 0.0000 | 0.0014 | 2.00 | 7,627 | **0.046** | 0.200 |
| R Pars Orbitalis | 0.0000 | -0.0005 | 0.0006 | 0.09 | 8,181 | 0.925 | 0.925 |
| L Superior Frontal Gyrus | 0.0000 | -0.0001 | 0.0001 | 0.40 | 3,626 | 0.687 | 0.862 |
| R Superior Frontal Gyrus | 0.0001 | 0.0000 | 0.0001 | 1.10 | 2,828 | 0.273 | 0.591 |
| L Precuneus | 0.0002 | 0.0000 | 0.0004 | 1.89 | 6,687 | 0.059 | 0.219 |
| R Precuneus | 0.0001 | 0.0000 | 0.0003 | 1.48 | 4,849 | 0.140 | 0.363 |

### **S Table 8.** Linear mixed effect model results for relationship between enrichment and rs-fMRI ROI at Baseline

| ROI - Outcome | 𝛽 | Lower | Upper | t | df | p | p adj |
| --- | --- | --- | --- | --- | --- | --- | --- |
| L Hippocampus | 0.009 | -0.005 | 0.022 | 1.27 | 7,702 | 0.204 | 0.883 |
| R Hippocampus | 0.009 | -0.004 | 0.022 | 1.37 | 7,897 | 0.171 | 0.883 |
| L Amygdala | -0.001 | -0.043 | 0.040 | -0.07 | 7,936 | 0.947 | 0.989 |
| R Amygdala | 0.013 | -0.039 | 0.064 | 0.49 | 7,928 | 0.627 | 0.983 |
| L Entorhinal Cortex | 0.044 | -0.363 | 0.452 | 0.21 | 7,922 | 0.831 | 0.989 |
| R Entorhinal Cortex | 0.096 | -0.321 | 0.512 | 0.45 | 7,947 | 0.653 | 0.983 |
| L Parahippocampus | -0.015 | -0.045 | 0.015 | -0.99 | 7,948 | 0.321 | 0.983 |
| R Parahippocampus | -0.016 | -0.049 | 0.017 | -0.94 | 7,942 | 0.346 | 0.983 |
| L Lateral Orbitofrontal Cortex | 0.015 | -0.028 | 0.057 | 0.67 | 7,949 | 0.502 | 0.983 |
| R Lateral Orbitofrontal Cortex | 0.014 | -0.035 | 0.062 | 0.56 | 7,950 | 0.575 | 0.983 |
| L Medial Orbitofrontal Cortex | 0.161 | 0.000 | 0.321 | 1.97 | 7,950 | **0.049** | 0.642 |
| R Medial Orbitofrontal Cortex | 0.186 | 0.044 | 0.328 | 2.57 | 7,949 | **0.010** | 0.261 |
| L Caudal Middle Frontal Gyrus | -0.006 | -0.014 | 0.002 | -1.54 | 6,769 | 0.124 | 0.809 |
| R Caudal Middle Frontal Gyrus | -0.008 | -0.018 | 0.002 | -1.61 | 5,414 | 0.108 | 0.809 |
| L Rostral Middle Frontal Gyrus | 0.037 | -0.094 | 0.167 | 0.55 | 6,795 | 0.581 | 0.983 |
| R Rostral Middle Frontal Gyrus | 0.046 | -0.322 | 0.413 | 0.24 | 3,825 | 0.808 | 0.989 |
| L Pars Opercularis | 0.000 | -0.009 | 0.009 | -0.03 | 7,933 | 0.979 | 0.989 |
| R Pars Opercularis | 0.004 | -0.007 | 0.015 | 0.74 | 7,849 | 0.461 | 0.983 |
| L Pars Triangularis | 0.012 | -0.046 | 0.069 | 0.40 | 7,554 | 0.689 | 0.983 |
| R Pars Triangularis | 0.015 | -0.013 | 0.042 | 1.05 | 7,935 | 0.295 | 0.983 |
| L Pars Orbitalis | 0.077 | -0.150 | 0.304 | 0.66 | 7,898 | 0.506 | 0.983 |
| R Pars Orbitalis | 0.109 | -0.135 | 0.352 | 0.88 | 7,914 | 0.381 | 0.983 |
| L Superior Frontal Gyrus | 0.000 | -0.025 | 0.025 | -0.03 | 2,695 | 0.979 | 0.989 |
| R Superior Frontal Gyrus | 0.000 | -0.030 | 0.029 | -0.01 | 2,991 | 0.989 | 0.989 |
| L Precuneus | -0.001 | -0.012 | 0.011 | -0.11 | 6,908 | 0.916 | 0.989 |
| R Precuneus | 0.002 | -0.009 | 0.013 | 0.36 | 7,403 | 0.719 | 0.983 |

### **S Table 9.** Linear mixed effect model results for relationship between PSMT scores and rs-fMRI ROI at Baseline

| ROI - Predictor | 𝛽 | Lower | Upper | t | df | p | p adj |
| --- | --- | --- | --- | --- | --- | --- | --- |
| L Hippocampus | 6.93 | -9.15 | 23.01 | 0.84 | 8,268 | 0.398 | 0.576 |
| R Hippocampus | -2.78 | -19.46 | 13.90 | -0.33 | 7,416 | 0.744 | 0.821 |
| L Amygdala | -0.35 | -5.72 | 5.01 | -0.13 | 6,401 | 0.897 | 0.897 |
| R Amygdala | -1.53 | -5.70 | 2.65 | -0.72 | 6,731 | 0.474 | 0.648 |
| L Entorhinal Cortex | -0.07 | -0.60 | 0.45 | -0.27 | 6,870 | 0.789 | 0.821 |
| R Entorhinal Cortex | -0.15 | -0.66 | 0.36 | -0.59 | 4,731 | 0.558 | 0.691 |
| L Parahippocampus | 3.78 | -3.21 | 10.77 | 1.06 | 5,616 | 0.289 | 0.575 |
| R Parahippocampus | -2.15 | -8.69 | 4.39 | -0.64 | 6,248 | 0.519 | 0.675 |
| L Lateral Orbitofrontal Cortex | 0.72 | -4.23 | 5.68 | 0.29 | 3,430 | 0.775 | 0.821 |
| R Lateral Orbitofrontal Cortex | 2.37 | -1.96 | 6.69 | 1.07 | 4,319 | 0.284 | 0.575 |
| L Medial Orbitofrontal Cortex | 0.63 | -0.73 | 1.99 | 0.90 | 5,095 | 0.366 | 0.575 |
| R Medial Orbitofrontal Cortex | 1.19 | -0.34 | 2.71 | 1.52 | 5,263 | 0.128 | 0.415 |
| L Caudal Middle Frontal Gyrus | 19.56 | -8.17 | 47.29 | 1.38 | 8,522 | 0.167 | 0.482 |
| R Caudal Middle Frontal Gyrus | 19.35 | -3.21 | 41.91 | 1.68 | 8,558 | 0.093 | 0.344 |
| L Rostral Middle Frontal Gyrus | 1.10 | -0.62 | 2.82 | 1.25 | 8,496 | 0.212 | 0.551 |
| R Rostral Middle Frontal Gyrus | 0.28 | -0.32 | 0.89 | 0.92 | 8,564 | 0.356 | 0.575 |
| L Pars Opercularis | 10.91 | -13.25 | 35.07 | 0.88 | 7,250 | 0.376 | 0.575 |
| R Pars Opercularis | 17.83 | -2.07 | 37.72 | 1.76 | 8,078 | 0.079 | 0.344 |
| L Pars Triangularis | 4.08 | 0.17 | 7.99 | 2.04 | 8,344 | **0.041** | 0.344 |
| R Pars Triangularis | 6.94 | -1.12 | 15.00 | 1.69 | 6,917 | 0.092 | 0.344 |
| L Pars Orbitalis | 0.14 | -0.82 | 1.10 | 0.29 | 7,232 | 0.773 | 0.821 |
| R Pars Orbitalis | 0.93 | 0.06 | 1.81 | 2.09 | 7,117 | **0.037** | 0.344 |
| L Superior Frontal Gyrus | 4.36 | -4.36 | 13.07 | 0.98 | 8,560 | 0.328 | 0.575 |
| R Superior Frontal Gyrus | 4.35 | -3.12 | 11.83 | 1.14 | 8,561 | 0.254 | 0.575 |
| L Precuneus | 23.68 | 4.58 | 42.78 | 2.43 | 8,496 | **0.015** | 0.344 |
| R Precuneus | 19.03 | -1.31 | 39.38 | 1.83 | 8,394 | 0.067 | 0.344 |

### **S Table 10.** Linear mixed effect model results for relationship between enrichment and tfMRI ROI at Baseline and Follow-up

| ROI - Outcome | 𝛽 | Lower | Upper | t | df | p | p adj |
| --- | --- | --- | --- | --- | --- | --- | --- |
| L Hippocampus | 0.06 | -0.24 | 0.36 | 0.41 | 2,552 | 0.682 | 0.762 |
| R Hippocampus | 0.04 | -0.27 | 0.35 | 0.24 | 2,969 | 0.814 | 0.814 |
| L Amygdala | 0.12 | -0.30 | 0.54 | 0.55 | 2,634 | 0.580 | 0.712 |
| R Amygdala | -0.32 | -0.76 | 0.11 | -1.46 | 2,730 | 0.144 | 0.267 |
| L Entorhinal Cortex | 0.18 | -0.92 | 1.27 | 0.32 | 2,578 | 0.753 | 0.783 |
| R Entorhinal Cortex | -0.39 | -1.84 | 1.07 | -0.52 | 1,768 | 0.602 | 0.712 |
| L Parahippocampus | -0.12 | -0.51 | 0.27 | -0.60 | 2,729 | 0.551 | 0.712 |
| R Parahippocampus | -0.19 | -0.61 | 0.23 | -0.89 | 2,550 | 0.375 | 0.573 |
| L Lateral Orbitofrontal Cortex | 0.21 | -0.36 | 0.78 | 0.72 | 2,944 | 0.472 | 0.681 |
| R Lateral Orbitofrontal Cortex | 0.42 | -0.20 | 1.05 | 1.33 | 2,861 | 0.184 | 0.299 |
| L Medial Orbitofrontal Cortex | 0.14 | -0.59 | 0.88 | 0.38 | 2,677 | 0.703 | 0.762 |
| R Medial Orbitofrontal Cortex | 0.19 | -0.52 | 0.90 | 0.53 | 6,421 | 0.594 | 0.712 |
| L Caudal Middle Frontal Gyrus | 0.52 | 0.24 | 0.79 | 3.71 | 2,830 | **<0.001** | **0.005** |
| R Caudal Middle Frontal Gyrus | 0.45 | 0.17 | 0.74 | 3.09 | 2,879 | **0.002** | **0.014** |
| L Rostral Middle Frontal Gyrus | 0.58 | 0.20 | 0.96 | 2.97 | 3,807 | **0.003** | **0.016** |
| R Rostral Middle Frontal Gyrus | 0.59 | 0.19 | 0.99 | 2.87 | 3,738 | **0.004** | **0.018** |
| L Pars Opercularis | 0.43 | 0.12 | 0.74 | 2.72 | 3,718 | **0.006** | **0.022** |
| R Pars Opercularis | 0.43 | 0.12 | 0.74 | 2.71 | 3,720 | **0.007** | **0.022** |
| L Pars Triangularis | 0.44 | 0.07 | 0.80 | 2.36 | 2,723 | **0.018** | 0.053 |
| R Pars Triangularis | 0.37 | 0.00 | 0.74 | 1.94 | 2,451 | 0.053 | 0.115 |
| L Pars Orbitalis | 0.60 | -0.25 | 1.46 | 1.38 | 3,002 | 0.166 | 0.288 |
| R Pars Orbitalis | 0.75 | -0.11 | 1.61 | 1.70 | 3,013 | 0.089 | 0.177 |
| L Superior Frontal Gyrus | 0.29 | 0.03 | 0.54 | 2.22 | 2,617 | **0.027** | 0.068 |
| R Superior Frontal Gyrus | 0.29 | 0.03 | 0.55 | 2.19 | 2,883 | **0.029** | 0.068 |
| L Precuneus | 0.48 | 0.20 | 0.75 | 3.40 | 2,627 | **0.001** | **0.009** |
| R Precuneus | 0.43 | 0.16 | 0.71 | 3.08 | 3,137 | **0.002** | **0.014** |

### **S Table 11.** Linear mixed effect model results for relationship between PSMT scores and tfMRI ROI at Baseline and Follow-up

| Predictor | 𝛽 | Lower | Upper | t | df | p | p adj |
| --- | --- | --- | --- | --- | --- | --- | --- |
| L Hippocampus | -0.41 | -1.29 | 0.46 | -0.93 | 6,620 | 0.353 | 0.540 |
| R Hippocampus | -0.27 | -1.11 | 0.58 | -0.61 | 6,620 | 0.539 | 0.667 |
| L Amygdala | -0.34 | -0.96 | 0.27 | -1.08 | 6,620 | 0.278 | 0.536 |
| R Amygdala | -0.09 | -0.69 | 0.52 | -0.28 | 6,620 | 0.780 | 0.865 |
| L Entorhinal Cortex | -0.02 | -0.27 | 0.22 | -0.17 | 6,620 | 0.865 | 0.865 |
| R Entorhinal Cortex | 0.13 | -0.05 | 0.31 | 1.40 | 6,619 | 0.161 | 0.349 |
| L Parahippocampus | -0.35 | -1.02 | 0.32 | -1.03 | 6,620 | 0.303 | 0.536 |
| R Parahippocampus | -0.07 | -0.69 | 0.55 | -0.23 | 6,620 | 0.819 | 0.865 |
| L Lateral Orbitofrontal Cortex | 0.24 | -0.23 | 0.70 | 1.01 | 6,619 | 0.312 | 0.536 |
| R Lateral Orbitofrontal Cortex | 0.05 | -0.38 | 0.47 | 0.21 | 6,620 | 0.832 | 0.865 |
| L Medial Orbitofrontal Cortex | -0.14 | -0.50 | 0.22 | -0.77 | 6,620 | 0.442 | 0.574 |
| R Medial Orbitofrontal Cortex | -0.17 | -0.53 | 0.20 | -0.89 | 6,617 | 0.376 | 0.543 |
| L Caudal Middle Frontal Gyrus | 2.36 | 1.40 | 3.32 | 4.83 | 6,619 | **<0.001** | **<0.001** |
| R Caudal Middle Frontal Gyrus | 1.42 | 0.52 | 2.33 | 3.09 | 6,620 | **0.002** | **0.018** |
| L Rostral Middle Frontal Gyrus | 1.01 | 0.32 | 1.71 | 2.86 | 6,617 | **0.004** | **0.025** |
| R Rostral Middle Frontal Gyrus | 0.95 | 0.29 | 1.60 | 2.82 | 6,614 | **0.005** | **0.025** |
| L Pars Opercularis | 1.43 | 0.57 | 2.29 | 3.26 | 6,614 | **0.001** | **0.015** |
| R Pars Opercularis | 0.75 | -0.10 | 1.60 | 1.72 | 6,616 | 0.086 | 0.279 |
| L Pars Triangularis | 0.59 | -0.13 | 1.31 | 1.61 | 6,620 | 0.107 | 0.279 |
| R Pars Triangularis | -0.17 | -0.87 | 0.53 | -0.48 | 6,620 | 0.631 | 0.746 |
| L Pars Orbitalis | 0.13 | -0.18 | 0.44 | 0.84 | 6,620 | 0.401 | 0.548 |
| R Pars Orbitalis | 0.15 | -0.16 | 0.46 | 0.97 | 6,620 | 0.330 | 0.536 |
| L Superior Frontal Gyrus | 1.01 | -0.02 | 2.04 | 1.92 | 6,620 | 0.055 | 0.206 |
| R Superior Frontal Gyrus | 0.85 | -0.16 | 1.85 | 1.65 | 6,620 | 0.099 | 0.279 |
| L Precuneus | 0.73 | -0.24 | 1.69 | 1.48 | 6,620 | 0.139 | 0.329 |
| R Precuneus | 1.19 | 0.22 | 2.15 | 2.41 | 6,620 | **0.016** | 0.069 |

### **S Table 12.** Linear mixed effect model results for relationship between enrichment and ROI volume at Baseline and Follow-up

| ROI - Outcome | 𝛽 | Lower | Upper | t | df | p | p adj |
| --- | --- | --- | --- | --- | --- | --- | --- |
| L Hippocampus | 87.86 | -180.67 | 356.38 | 0.64 | 7,374 | 0.521 | 0.713 |
| R Hippocampus | 256.02 | -30.49 | 542.54 | 1.75 | 6,885 | 0.080 | 0.173 |
| L Amygdala | 207.75 | 45.18 | 370.31 | 2.50 | 7,545 | **0.012** | **0.047** |
| R Amygdala | 167.15 | 16.73 | 317.57 | 2.18 | 7,073 | **0.029** | 0.089 |
| L Entorhinal Cortex | 292.95 | -52.77 | 638.66 | 1.66 | 7,394 | 0.097 | 0.194 |
| R Entorhinal Cortex | 424.86 | 91.06 | 758.66 | 2.49 | 5,015 | **0.013** | **0.047** |
| L Parahippocampus | 380.38 | 86.33 | 674.43 | 2.54 | 7,457 | **0.011** | **0.047** |
| R Parahippocampus | 245.52 | 2.98 | 488.05 | 1.98 | 6,633 | **0.047** | 0.123 |
| L Lateral Orbitofrontal Cortex | 743.98 | 69.34 | 1,418.62 | 2.16 | 7,571 | **0.031** | 0.089 |
| R Lateral Orbitofrontal Cortex | 638.22 | -54.81 | 1,331.24 | 1.80 | 7,563 | 0.071 | 0.168 |
| L Medial Orbitofrontal Cortex | -16.68 | -550.27 | 516.92 | -0.06 | 7,560 | 0.951 | 0.951 |
| R Medial Orbitofrontal Cortex | 46.81 | -497.76 | 591.39 | 0.17 | 7,570 | 0.866 | 0.907 |
| L Caudal Middle Frontal Gyrus | 845.60 | -302.26 | 1,993.46 | 1.44 | 7,571 | 0.149 | 0.272 |
| R Caudal Middle Frontal Gyrus | -157.16 | -1,319.02 | 1,004.71 | -0.27 | 7,570 | 0.791 | 0.907 |
| L Rostral Middle Frontal Gyrus | 2,944.78 | 1,073.63 | 4,815.92 | 3.08 | 7,565 | **0.002** | **0.018** |
| R Rostral Middle Frontal Gyrus | 3,697.95 | 1,636.03 | 5,759.88 | 3.52 | 7,572 | **<0.001** | **0.006** |
| L Pars Opercularis | -228.47 | -1,075.18 | 618.24 | -0.53 | 7,445 | 0.597 | 0.776 |
| R Pars Opercularis | 75.19 | -606.35 | 756.73 | 0.22 | 7,480 | 0.829 | 0.907 |
| L Pars Triangularis | -251.44 | -933.60 | 430.73 | -0.72 | 7,348 | 0.470 | 0.679 |
| R Pars Triangularis | -558.52 | -1,351.81 | 234.77 | -1.38 | 7,447 | 0.168 | 0.272 |
| L Pars Orbitalis | 496.34 | 170.94 | 821.73 | 2.99 | 7,544 | **0.003** | **0.018** |
| R Pars Orbitalis | 701.63 | 312.16 | 1,091.10 | 3.53 | 7,498 | **<0.001** | **0.006** |
| L Superior Frontal Gyrus | -320.06 | -2,601.78 | 1,961.65 | -0.27 | 7,571 | 0.783 | 0.907 |
| R Superior Frontal Gyrus | 188.66 | -2,116.57 | 2,493.90 | 0.16 | 7,569 | 0.873 | 0.907 |
| L Precuneus | 589.28 | -514.48 | 1,693.05 | 1.05 | 7,561 | 0.295 | 0.452 |
| R Precuneus | 795.03 | -327.37 | 1,917.42 | 1.39 | 7,572 | 0.165 | 0.272 |

### **S Table 13.** Linear mixed effect model results for relationship between PSMT scores and ROI volume at Baseline and Follow-up

| ROI - Predictor | 𝛽 | Lower | Upper | t | df | p | p adj |
| --- | --- | --- | --- | --- | --- | --- | --- |
| L Hippocampus | 0.0008 | 0.0000 | 0.0016 | 1.87 | 7,357 | 0.062 | 0.403 |
| R Hippocampus | 0.0008 | 0.0000 | 0.0016 | 1.92 | 7,685 | 0.055 | 0.403 |
| L Amygdala | 0.0000 | -0.0014 | 0.0014 | 0.00 | 6,452 | 1.000 | 1.000 |
| R Amygdala | -0.0007 | -0.0022 | 0.0008 | -0.91 | 7,647 | 0.363 | 0.702 |
| L Entorhinal Cortex | 0.0010 | 0.0003 | 0.0017 | 3.00 | 7,345 | **0.003** | 0.070 |
| R Entorhinal Cortex | 0.0000 | -0.0007 | 0.0006 | -0.09 | 7,797 | 0.931 | 0.968 |
| L Parahippocampus | 0.0002 | -0.0005 | 0.0010 | 0.61 | 7,143 | 0.543 | 0.743 |
| R Parahippocampus | -0.0005 | -0.0015 | 0.0004 | -1.11 | 7,738 | 0.268 | 0.698 |
| L Lateral Orbitofrontal Cortex | 0.0003 | 0.0000 | 0.0006 | 1.73 | 4,085 | 0.083 | 0.431 |
| R Lateral Orbitofrontal Cortex | 0.0002 | -0.0001 | 0.0005 | 1.14 | 763 | 0.254 | 0.698 |
| L Medial Orbitofrontal Cortex | 0.0001 | -0.0003 | 0.0005 | 0.44 | 532 | 0.660 | 0.806 |
| R Medial Orbitofrontal Cortex | -0.0001 | -0.0005 | 0.0003 | -0.37 | 2,420 | 0.713 | 0.806 |
| L Caudal Middle Frontal Gyrus | 0.0002 | 0.0000 | 0.0004 | 1.56 | 4,280 | 0.119 | 0.515 |
| R Caudal Middle Frontal Gyrus | 0.0001 | -0.0001 | 0.0003 | 1.18 | 4,571 | 0.239 | 0.698 |
| L Rostral Middle Frontal Gyrus | -0.0001 | -0.0002 | 0.0001 | -0.91 | 5,360 | 0.364 | 0.702 |
| R Rostral Middle Frontal Gyrus | 0.0001 | -0.0001 | 0.0002 | 0.98 | 4,165 | 0.327 | 0.702 |
| L Pars Opercularis | 0.0000 | -0.0002 | 0.0003 | 0.30 | 7,261 | 0.765 | 0.829 |
| R Pars Opercularis | 0.0004 | 0.0001 | 0.0007 | 2.42 | 7,062 | **0.015** | 0.199 |
| L Pars Triangularis | 0.0001 | -0.0002 | 0.0005 | 0.83 | 7,407 | 0.404 | 0.702 |
| R Pars Triangularis | 0.0001 | -0.0002 | 0.0004 | 0.83 | 7,248 | 0.408 | 0.702 |
| L Pars Orbitalis | -0.0003 | -0.0010 | 0.0004 | -0.79 | 6,443 | 0.432 | 0.702 |
| R Pars Orbitalis | 0.0001 | -0.0005 | 0.0007 | 0.45 | 6,966 | 0.654 | 0.806 |
| L Superior Frontal Gyrus | 0.0001 | 0.0000 | 0.0002 | 1.39 | 2,573 | 0.163 | 0.607 |
| R Superior Frontal Gyrus | 0.0000 | -0.0001 | 0.0001 | 0.73 | 2,133 | 0.465 | 0.711 |
| L Precuneus | 0.0001 | -0.0001 | 0.0003 | 0.65 | 5,523 | 0.519 | 0.743 |
| R Precuneus | 0.0000 | -0.0002 | 0.0002 | 0.38 | 3,789 | 0.705 | 0.806 |

### **S Table 14.** Linear mixed effect model results for relationship between enrichment and rs-fMRI ROI at Baseline and Follow-up

| ROI - Outcome | 𝛽 | Lower | Upper | t | df | p | p adj |
| --- | --- | --- | --- | --- | --- | --- | --- |
| L Hippocampus | 0.009 | -0.005 | 0.024 | 1.27 | 7,023 | 0.202 | 0.744 |
| R Hippocampus | 0.011 | -0.003 | 0.025 | 1.56 | 7,226 | 0.120 | 0.744 |
| L Amygdala | -0.006 | -0.043 | 0.031 | -0.31 | 7,278 | 0.758 | 0.910 |
| R Amygdala | 0.006 | -0.048 | 0.061 | 0.23 | 7,257 | 0.819 | 0.910 |
| L Entorhinal Cortex | 0.064 | -0.304 | 0.433 | 0.34 | 7,272 | 0.732 | 0.910 |
| R Entorhinal Cortex | 0.080 | -0.349 | 0.509 | 0.36 | 7,280 | 0.716 | 0.910 |
| L Parahippocampus | -0.018 | -0.050 | 0.013 | -1.14 | 7,282 | 0.255 | 0.744 |
| R Parahippocampus | -0.017 | -0.052 | 0.019 | -0.92 | 7,274 | 0.355 | 0.840 |
| L Lateral Orbitofrontal Cortex | 0.005 | -0.041 | 0.050 | 0.20 | 7,283 | 0.838 | 0.910 |
| R Lateral Orbitofrontal Cortex | 0.006 | -0.046 | 0.058 | 0.23 | 7,283 | 0.820 | 0.910 |
| L Medial Orbitofrontal Cortex | 0.129 | -0.040 | 0.299 | 1.49 | 7,282 | 0.136 | 0.744 |
| R Medial Orbitofrontal Cortex | 0.160 | 0.008 | 0.312 | 2.07 | 7,280 | **0.039** | 0.744 |
| L Caudal Middle Frontal Gyrus | -0.006 | -0.014 | 0.003 | -1.29 | 6,140 | 0.196 | 0.744 |
| R Caudal Middle Frontal Gyrus | -0.008 | -0.019 | 0.002 | -1.57 | 4,758 | 0.116 | 0.744 |
| L Rostral Middle Frontal Gyrus | 0.035 | -0.107 | 0.178 | 0.48 | 6,210 | 0.628 | 0.910 |
| R Rostral Middle Frontal Gyrus | 0.043 | -0.361 | 0.446 | 0.21 | 3,707 | 0.836 | 0.910 |
| L Pars Opercularis | 0.001 | -0.008 | 0.011 | 0.27 | 7,267 | 0.785 | 0.910 |
| R Pars Opercularis | 0.007 | -0.005 | 0.018 | 1.13 | 7,174 | 0.258 | 0.744 |
| L Pars Triangularis | 0.013 | -0.050 | 0.075 | 0.40 | 6,837 | 0.693 | 0.910 |
| R Pars Triangularis | 0.019 | -0.011 | 0.048 | 1.24 | 7,262 | 0.216 | 0.744 |
| L Pars Orbitalis | 0.097 | -0.133 | 0.328 | 0.83 | 7,235 | 0.409 | 0.886 |
| R Pars Orbitalis | 0.139 | -0.120 | 0.398 | 1.05 | 7,241 | 0.293 | 0.761 |
| L Superior Frontal Gyrus | 0.000 | -0.027 | 0.028 | 0.02 | 2,635 | 0.982 | 0.982 |
| R Superior Frontal Gyrus | 0.001 | -0.032 | 0.033 | 0.04 | 2,943 | 0.971 | 0.982 |
| L Precuneus | -0.002 | -0.014 | 0.010 | -0.30 | 6,406 | 0.767 | 0.910 |
| R Precuneus | 0.001 | -0.010 | 0.012 | 0.20 | 6,801 | 0.840 | 0.910 |

### **S Table 15.** Linear mixed effect model results for relationship between PSMT scores and rs-fMRI ROI at Baseline and Follow-up

| ROI - Predictor | 𝛽 | Lower | Upper | t | df | p | p adj |
| --- | --- | --- | --- | --- | --- | --- | --- |
| L Hippocampus | 4.89 | -11.63 | 21.41 | 0.58 | 7,268 | 0.562 | 0.995 |
| R Hippocampus | -6.73 | -23.80 | 10.35 | -0.77 | 6,419 | 0.440 | 0.995 |
| L Amygdala | -0.25 | -6.53 | 6.03 | -0.08 | 4,745 | 0.937 | 0.995 |
| R Amygdala | 2.29 | -1.91 | 6.48 | 1.07 | 5,630 | 0.285 | 0.995 |
| L Entorhinal Cortex | 0.06 | -0.56 | 0.68 | 0.20 | 5,204 | 0.845 | 0.995 |
| R Entorhinal Cortex | -0.18 | -0.71 | 0.35 | -0.66 | 3,802 | 0.510 | 0.995 |
| L Parahippocampus | -4.64 | -11.71 | 2.44 | -1.28 | 4,790 | 0.199 | 0.864 |
| R Parahippocampus | -5.61 | -12.20 | 0.97 | -1.67 | 5,387 | 0.095 | 0.823 |
| L Lateral Orbitofrontal Cortex | 3.56 | -1.52 | 8.65 | 1.37 | 3,190 | 0.170 | 0.864 |
| R Lateral Orbitofrontal Cortex | -0.36 | -4.71 | 3.99 | -0.16 | 4,000 | 0.871 | 0.995 |
| L Medial Orbitofrontal Cortex | 0.21 | -1.17 | 1.60 | 0.30 | 4,773 | 0.762 | 0.995 |
| R Medial Orbitofrontal Cortex | 0.56 | -0.98 | 2.10 | 0.71 | 4,971 | 0.476 | 0.995 |
| L Caudal Middle Frontal Gyrus | 6.56 | -21.64 | 34.76 | 0.46 | 7,483 | 0.648 | 0.995 |
| R Caudal Middle Frontal Gyrus | 0.13 | -22.36 | 22.62 | 0.01 | 7,503 | 0.991 | 0.995 |
| L Rostral Middle Frontal Gyrus | 0.08 | -1.60 | 1.76 | 0.09 | 7,442 | 0.928 | 0.995 |
| R Rostral Middle Frontal Gyrus | 0.00 | -0.58 | 0.59 | 0.01 | 7,507 | 0.993 | 0.995 |
| L Pars Opercularis | -25.17 | -50.32 | -0.01 | -1.96 | 6,221 | **0.050** | 0.649 |
| R Pars Opercularis | -3.53 | -23.95 | 16.89 | -0.34 | 7,112 | 0.735 | 0.995 |
| L Pars Triangularis | 1.63 | -2.18 | 5.44 | 0.84 | 7,345 | 0.402 | 0.995 |
| R Pars Triangularis | 0.84 | -7.13 | 8.82 | 0.21 | 6,217 | 0.836 | 0.995 |
| L Pars Orbitalis | 0.00 | -1.00 | 1.01 | 0.01 | 6,093 | 0.995 | 0.995 |
| R Pars Orbitalis | 0.16 | -0.74 | 1.05 | 0.35 | 6,235 | 0.730 | 0.995 |
| L Superior Frontal Gyrus | -0.34 | -8.79 | 8.12 | -0.08 | 7,503 | 0.938 | 0.995 |
| R Superior Frontal Gyrus | -0.32 | -7.57 | 6.93 | -0.09 | 7,504 | 0.931 | 0.995 |
| L Precuneus | 22.12 | 2.26 | 41.97 | 2.18 | 7,407 | **0.029** | 0.649 |
| R Precuneus | 16.38 | -4.67 | 37.44 | 1.53 | 7,293 | 0.127 | 0.827 |

### **S Table 16.** Linear mixed effect model results for relationship between enrichment and tfMRI ROI at Follow-up

| ROI - Outcome | 𝛽 | Lower | Upper | t | df | p | p adj |
| --- | --- | --- | --- | --- | --- | --- | --- |
| L Hippocampus | -0.29 | -0.59 | 0.01 | -1.87 | 1,882 | 0.062 | 0.396 |
| R Hippocampus | -0.26 | -0.59 | 0.06 | -1.58 | 4,957 | 0.114 | 0.396 |
| L Amygdala | -0.17 | -0.61 | 0.28 | -0.74 | 4,957 | 0.461 | 0.705 |
| R Amygdala | -0.35 | -0.76 | 0.06 | -1.67 | 4,957 | 0.095 | 0.396 |
| L Entorhinal Cortex | -1.05 | -2.05 | -0.04 | -2.04 | 4,957 | **0.041** | 0.396 |
| R Entorhinal Cortex | -0.71 | -1.71 | 0.29 | -1.40 | 2,797 | 0.162 | 0.397 |
| L Parahippocampus | -0.05 | -0.44 | 0.35 | -0.25 | 4,957 | 0.806 | 0.935 |
| R Parahippocampus | 0.03 | -0.38 | 0.43 | 0.13 | 2,776 | 0.899 | 0.935 |
| L Lateral Orbitofrontal Cortex | -0.60 | -1.24 | 0.04 | -1.84 | 4,957 | 0.065 | 0.396 |
| R Lateral Orbitofrontal Cortex | -0.40 | -0.95 | 0.16 | -1.41 | 2,463 | 0.160 | 0.397 |
| L Medial Orbitofrontal Cortex | -0.66 | -1.46 | 0.15 | -1.59 | 2,927 | 0.111 | 0.396 |
| R Medial Orbitofrontal Cortex | -0.74 | -1.47 | -0.01 | -1.98 | 2,911 | **0.047** | 0.396 |
| L Caudal Middle Frontal Gyrus | 0.17 | -0.15 | 0.50 | 1.03 | 3,237 | 0.303 | 0.562 |
| R Caudal Middle Frontal Gyrus | 0.16 | -0.17 | 0.48 | 0.94 | 4,957 | 0.347 | 0.602 |
| L Rostral Middle Frontal Gyrus | 0.01 | -0.38 | 0.41 | 0.07 | 3,297 | 0.945 | 0.945 |
| R Rostral Middle Frontal Gyrus | 0.03 | -0.38 | 0.44 | 0.14 | 3,280 | 0.885 | 0.935 |
| L Pars Opercularis | 0.06 | -0.27 | 0.39 | 0.34 | 3,512 | 0.731 | 0.935 |
| R Pars Opercularis | 0.05 | -0.29 | 0.39 | 0.29 | 3,573 | 0.772 | 0.935 |
| L Pars Triangularis | -0.10 | -0.50 | 0.31 | -0.47 | 4,957 | 0.641 | 0.926 |
| R Pars Triangularis | -0.33 | -0.82 | 0.16 | -1.33 | 4,957 | 0.183 | 0.397 |
| L Pars Orbitalis | -0.52 | -1.27 | 0.23 | -1.35 | 4,957 | 0.177 | 0.397 |
| R Pars Orbitalis | -0.56 | -1.27 | 0.15 | -1.55 | 2,588 | 0.122 | 0.396 |
| L Superior Frontal Gyrus | -0.16 | -0.45 | 0.12 | -1.11 | 3,130 | 0.269 | 0.538 |
| R Superior Frontal Gyrus | -0.12 | -0.40 | 0.15 | -0.89 | 3,260 | 0.372 | 0.604 |
| L Precuneus | 0.05 | -0.27 | 0.36 | 0.29 | 2,984 | 0.774 | 0.935 |
| R Precuneus | 0.03 | -0.28 | 0.34 | 0.18 | 4,957 | 0.857 | 0.935 |

### **S Table 17.** Linear mixed effect model results for relationship between PSMT scores and tfMRI ROI at Follow-up

| ROI - Predictor | 𝛽 | Lower | Upper | t | df | p | p adj |
| --- | --- | --- | --- | --- | --- | --- | --- |
| L Hippocampus | -2.03 | -3.22 | -0.84 | -3.34 | 5,155 | **0.001** | **0.004** |
| R Hippocampus | -0.20 | -1.29 | 0.89 | -0.35 | 5,155 | 0.723 | 0.777 |
| L Amygdala | -0.31 | -1.11 | 0.49 | -0.75 | 5,152 | 0.454 | 0.726 |
| R Amygdala | 0.19 | -0.67 | 1.06 | 0.44 | 5,156 | 0.663 | 0.777 |
| L Entorhinal Cortex | -0.26 | -0.61 | 0.10 | -1.41 | 5,144 | 0.160 | 0.348 |
| R Entorhinal Cortex | -0.10 | -0.46 | 0.25 | -0.57 | 5,155 | 0.568 | 0.726 |
| L Parahippocampus | -0.51 | -1.42 | 0.40 | -1.10 | 5,156 | 0.270 | 0.539 |
| R Parahippocampus | -0.24 | -1.12 | 0.63 | -0.54 | 5,156 | 0.586 | 0.726 |
| L Lateral Orbitofrontal Cortex | -0.09 | -0.66 | 0.47 | -0.33 | 5,154 | 0.741 | 0.777 |
| R Lateral Orbitofrontal Cortex | 0.19 | -0.47 | 0.84 | 0.56 | 5,155 | 0.577 | 0.726 |
| L Medial Orbitofrontal Cortex | -0.41 | -0.85 | 0.04 | -1.79 | 5,156 | 0.073 | 0.211 |
| R Medial Orbitofrontal Cortex | -0.35 | -0.84 | 0.14 | -1.40 | 5,155 | 0.160 | 0.348 |
| L Caudal Middle Frontal Gyrus | 3.27 | 2.18 | 4.36 | 5.89 | 5,155 | **<0.001** | **<0.001** |
| R Caudal Middle Frontal Gyrus | 3.43 | 2.34 | 4.51 | 6.19 | 5,155 | **<0.001** | **<0.001** |
| L Rostral Middle Frontal Gyrus | 1.78 | 0.87 | 2.68 | 3.85 | 5,156 | **<0.001** | **0.001** |
| R Rostral Middle Frontal Gyrus | 1.80 | 0.94 | 2.65 | 4.10 | 5,156 | **<0.001** | **<0.001** |
| L Pars Opercularis | 1.83 | 0.75 | 2.91 | 3.32 | 5,150 | **0.001** | **0.004** |
| R Pars Opercularis | 1.08 | 0.03 | 2.13 | 2.02 | 5,147 | **0.044** | 0.142 |
| L Pars Triangularis | 0.07 | -0.80 | 0.93 | 0.15 | 5,155 | 0.882 | 0.882 |
| R Pars Triangularis | -0.28 | -1.01 | 0.45 | -0.75 | 5,155 | 0.454 | 0.726 |
| L Pars Orbitalis | -0.08 | -0.56 | 0.40 | -0.32 | 5,147 | 0.747 | 0.777 |
| R Pars Orbitalis | 0.15 | -0.35 | 0.66 | 0.60 | 5,155 | 0.552 | 0.726 |
| L Superior Frontal Gyrus | 0.54 | -0.71 | 1.80 | 0.85 | 5,155 | 0.396 | 0.726 |
| R Superior Frontal Gyrus | 1.02 | -0.29 | 2.33 | 1.53 | 5,156 | 0.126 | 0.328 |
| L Precuneus | 0.38 | -0.72 | 1.49 | 0.68 | 5,156 | 0.496 | 0.726 |
| R Precuneus | 1.26 | 0.15 | 2.37 | 2.22 | 5,152 | **0.026** | 0.098 |

### **S Table 18.** Linear mixed effect model results for relationship between enrichment and ROI volume at Follow-up

| ROI - Outcome | 𝛽 | Lower | Upper | t | df | p | p adj |
| --- | --- | --- | --- | --- | --- | --- | --- |
| L Hippocampus | 96.37 | -221.96 | 414.69 | 0.59 | 5,004 | 0.553 | 0.757 |
| R Hippocampus | 250.96 | -87.25 | 589.17 | 1.45 | 4,564 | 0.146 | 0.292 |
| L Amygdala | 260.44 | 66.78 | 454.10 | 2.64 | 5,492 | **0.008** | **0.044** |
| R Amygdala | 290.48 | 114.07 | 466.88 | 3.23 | 5,175 | **0.001** | **0.011** |
| L Entorhinal Cortex | 412.55 | 4.83 | 820.26 | 1.98 | 5,296 | **0.047** | 0.112 |
| R Entorhinal Cortex | 654.71 | 280.32 | 1,029.10 | 3.43 | 3,141 | **0.001** | **0.008** |
| L Parahippocampus | 441.00 | 96.11 | 785.90 | 2.51 | 5,381 | **0.012** | **0.050** |
| R Parahippocampus | 176.53 | -109.20 | 462.25 | 1.21 | 4,763 | 0.226 | 0.385 |
| L Lateral Orbitofrontal Cortex | 805.64 | 12.20 | 1,599.08 | 1.99 | 5,498 | **0.047** | 0.112 |
| R Lateral Orbitofrontal Cortex | 900.58 | 86.48 | 1,714.69 | 2.17 | 5,497 | **0.030** | 0.087 |
| L Medial Orbitofrontal Cortex | 89.24 | -531.24 | 709.71 | 0.28 | 5,495 | 0.778 | 0.810 |
| R Medial Orbitofrontal Cortex | -137.19 | -766.08 | 491.70 | -0.43 | 5,504 | 0.669 | 0.796 |
| L Caudal Middle Frontal Gyrus | 468.46 | -865.12 | 1,802.04 | 0.69 | 5,502 | 0.491 | 0.751 |
| R Caudal Middle Frontal Gyrus | -292.41 | -1,634.97 | 1,050.14 | -0.43 | 5,501 | 0.669 | 0.796 |
| L Rostral Middle Frontal Gyrus | 2,427.14 | 271.84 | 4,582.43 | 2.21 | 5,490 | **0.027** | 0.087 |
| R Rostral Middle Frontal Gyrus | 4,813.18 | 2,419.09 | 7,207.27 | 3.94 | 5,503 | **<0.001** | **0.002** |
| L Pars Opercularis | -212.61 | -1,201.49 | 776.26 | -0.42 | 5,224 | 0.673 | 0.796 |
| R Pars Opercularis | 101.31 | -678.50 | 881.12 | 0.25 | 5,315 | 0.799 | 0.810 |
| L Pars Triangularis | -622.95 | -1,403.73 | 157.83 | -1.56 | 5,222 | 0.118 | 0.256 |
| R Pars Triangularis | -549.88 | -1,460.56 | 360.80 | -1.18 | 5,357 | 0.237 | 0.385 |
| L Pars Orbitalis | 473.94 | 98.65 | 849.24 | 2.48 | 5,403 | **0.013** | **0.050** |
| R Pars Orbitalis | 654.81 | 201.80 | 1,107.82 | 2.83 | 5,347 | **0.005** | **0.030** |
| L Superior Frontal Gyrus | -1,636.78 | -4,312.02 | 1,038.45 | -1.20 | 5,505 | 0.231 | 0.385 |
| R Superior Frontal Gyrus | 331.14 | -2,366.98 | 3,029.26 | 0.24 | 5,504 | 0.810 | 0.810 |
| L Precuneus | 398.33 | -866.58 | 1,663.24 | 0.62 | 5,469 | 0.537 | 0.757 |
| R Precuneus | 235.98 | -1,040.89 | 1,512.85 | 0.36 | 5,501 | 0.717 | 0.810 |

### **S Table 19.** Linear mixed effect model results for relationship between PSMT scores and ROI volume at Follow-up

| ROI - Predictor | 𝛽 | Lower | Upper | t | df | p | p adj |
| --- | --- | --- | --- | --- | --- | --- | --- |
| L Hippocampus | 0.0011 | 0.0000 | 0.0021 | 2.02 | 5,587 | **0.043** | 0.377 |
| R Hippocampus | 0.0012 | 0.0002 | 0.0022 | 2.39 | 5,678 | **0.017** | 0.250 |
| L Amygdala | -0.0001 | -0.0018 | 0.0016 | -0.10 | 4,038 | 0.921 | 0.941 |
| R Amygdala | -0.0002 | -0.0021 | 0.0016 | -0.23 | 5,481 | 0.815 | 0.941 |
| L Entorhinal Cortex | 0.0010 | 0.0002 | 0.0018 | 2.34 | 5,298 | **0.019** | 0.250 |
| R Entorhinal Cortex | 0.0002 | -0.0006 | 0.0011 | 0.48 | 5,723 | 0.630 | 0.883 |
| L Parahippocampus | 0.0004 | -0.0006 | 0.0013 | 0.75 | 5,230 | 0.455 | 0.844 |
| R Parahippocampus | -0.0007 | -0.0018 | 0.0005 | -1.11 | 5,681 | 0.267 | 0.844 |
| L Lateral Orbitofrontal Cortex | 0.0001 | -0.0003 | 0.0006 | 0.69 | 3,083 | 0.487 | 0.844 |
| R Lateral Orbitofrontal Cortex | 0.0001 | -0.0003 | 0.0004 | 0.35 | 523 | 0.728 | 0.941 |
| L Medial Orbitofrontal Cortex | -0.0001 | -0.0006 | 0.0003 | -0.55 | 435 | 0.586 | 0.883 |
| R Medial Orbitofrontal Cortex | -0.0002 | -0.0007 | 0.0003 | -0.90 | 1,792 | 0.368 | 0.844 |
| L Caudal Middle Frontal Gyrus | 0.0000 | -0.0003 | 0.0002 | -0.07 | 3,353 | 0.941 | 0.941 |
| R Caudal Middle Frontal Gyrus | 0.0000 | -0.0002 | 0.0003 | 0.18 | 3,383 | 0.854 | 0.941 |
| L Rostral Middle Frontal Gyrus | -0.0001 | -0.0002 | 0.0001 | -0.76 | 4,269 | 0.448 | 0.844 |
| R Rostral Middle Frontal Gyrus | 0.0000 | -0.0001 | 0.0002 | 0.46 | 3,143 | 0.645 | 0.883 |
| L Pars Opercularis | 0.0000 | -0.0004 | 0.0003 | -0.17 | 5,491 | 0.864 | 0.941 |
| R Pars Opercularis | 0.0002 | -0.0003 | 0.0006 | 0.77 | 5,394 | 0.438 | 0.844 |
| L Pars Triangularis | 0.0002 | -0.0002 | 0.0006 | 0.81 | 5,459 | 0.419 | 0.844 |
| R Pars Triangularis | 0.0002 | -0.0002 | 0.0005 | 0.92 | 5,314 | 0.360 | 0.844 |
| L Pars Orbitalis | -0.0003 | -0.0012 | 0.0005 | -0.78 | 5,206 | 0.437 | 0.844 |
| R Pars Orbitalis | -0.0003 | -0.0010 | 0.0004 | -0.82 | 5,352 | 0.411 | 0.844 |
| L Superior Frontal Gyrus | 0.0000 | -0.0001 | 0.0002 | 0.76 | 2,188 | 0.448 | 0.844 |
| R Superior Frontal Gyrus | 0.0000 | -0.0001 | 0.0001 | 0.12 | 1,783 | 0.905 | 0.941 |
| L Precuneus | 0.0001 | -0.0002 | 0.0003 | 0.56 | 4,223 | 0.574 | 0.883 |
| R Precuneus | 0.0001 | -0.0002 | 0.0004 | 0.75 | 2,986 | 0.454 | 0.844 |

### **S Table 20.** Linear mixed effect model results for relationship between enrichment and rs-fMRI ROI at Follow-up

| ROI - Outcome | 𝛽 | Lower | Upper | t | df | p | p adj |
| --- | --- | --- | --- | --- | --- | --- | --- |
| L Hippocampus | 0.012 | -0.010 | 0.034 | 1.09 | 5,379 | 0.275 | 0.632 |
| R Hippocampus | 0.016 | -0.004 | 0.036 | 1.60 | 5,389 | 0.109 | 0.404 |
| L Amygdala | 0.058 | -0.021 | 0.137 | 1.43 | 5,405 | 0.151 | 0.433 |
| R Amygdala | 0.029 | -0.042 | 0.099 | 0.80 | 5,405 | 0.424 | 0.688 |
| L Entorhinal Cortex | 0.297 | -0.274 | 0.868 | 1.02 | 5,403 | 0.308 | 0.632 |
| R Entorhinal Cortex | 0.575 | -0.196 | 1.345 | 1.46 | 5,403 | 0.144 | 0.433 |
| L Parahippocampus | -0.030 | -0.092 | 0.032 | -0.95 | 5,404 | 0.341 | 0.633 |
| R Parahippocampus | -0.005 | -0.063 | 0.053 | -0.17 | 5,405 | 0.868 | 0.903 |
| L Lateral Orbitofrontal Cortex | 0.002 | -0.080 | 0.083 | 0.05 | 5,405 | 0.964 | 0.964 |
| R Lateral Orbitofrontal Cortex | 0.052 | -0.001 | 0.105 | 1.93 | 5,399 | 0.054 | 0.350 |
| L Medial Orbitofrontal Cortex | 0.073 | -0.127 | 0.273 | 0.72 | 5,400 | 0.472 | 0.688 |
| R Medial Orbitofrontal Cortex | 0.056 | -0.135 | 0.247 | 0.58 | 5,403 | 0.563 | 0.688 |
| L Caudal Middle Frontal Gyrus | -0.011 | -0.020 | -0.003 | -2.54 | 5,284 | **0.011** | 0.275 |
| R Caudal Middle Frontal Gyrus | -0.007 | -0.016 | 0.001 | -1.67 | 5,281 | 0.096 | 0.404 |
| L Rostral Middle Frontal Gyrus | 0.009 | -0.024 | 0.042 | 0.54 | 5,403 | 0.588 | 0.688 |
| R Rostral Middle Frontal Gyrus | 0.018 | -0.049 | 0.084 | 0.52 | 5,393 | 0.602 | 0.688 |
| L Pars Opercularis | -0.006 | -0.017 | 0.005 | -1.00 | 5,405 | 0.316 | 0.632 |
| R Pars Opercularis | 0.005 | -0.009 | 0.019 | 0.66 | 5,390 | 0.509 | 0.688 |
| L Pars Triangularis | -0.026 | -0.058 | 0.005 | -1.63 | 5,405 | 0.104 | 0.404 |
| R Pars Triangularis | -0.005 | -0.025 | 0.015 | -0.51 | 5,401 | 0.609 | 0.688 |
| L Pars Orbitalis | -0.107 | -0.260 | 0.045 | -1.38 | 5,390 | 0.167 | 0.433 |
| R Pars Orbitalis | 0.049 | -0.103 | 0.201 | 0.63 | 5,403 | 0.531 | 0.688 |
| L Superior Frontal Gyrus | 0.001 | -0.004 | 0.007 | 0.46 | 5,393 | 0.645 | 0.698 |
| R Superior Frontal Gyrus | -0.002 | -0.008 | 0.004 | -0.69 | 5,401 | 0.492 | 0.688 |
| L Precuneus | -0.017 | -0.032 | -0.001 | -2.07 | 4,675 | **0.038** | 0.330 |
| R Precuneus | -0.017 | -0.032 | -0.003 | -2.31 | 4,682 | **0.021** | 0.275 |

### **S Table 21.** Linear mixed effect model results for relationship between PSMT scores and rs-fMRI ROI at Follow-up

| ROI - Predictor | 𝛽 | Lower | Upper | t | df | p | p adj |
| --- | --- | --- | --- | --- | --- | --- | --- |
| L Hippocampus | -13.52 | -29.04 | 2.01 | -1.71 | 4,302 | 0.088 | 0.558 |
| R Hippocampus | -7.13 | -24.32 | 10.06 | -0.81 | 3,493 | 0.416 | 0.760 |
| L Amygdala | -1.67 | -5.88 | 2.55 | -0.77 | 2,037 | 0.439 | 0.760 |
| R Amygdala | -0.78 | -5.48 | 3.91 | -0.33 | 1,429 | 0.744 | 0.921 |
| L Entorhinal Cortex | 0.19 | -0.36 | 0.74 | 0.67 | 1,168 | 0.502 | 0.760 |
| R Entorhinal Cortex | 0.00 | -0.42 | 0.42 | 0.00 | 1,035 | 0.999 | 0.999 |
| L Parahippocampus | 1.97 | -2.94 | 6.88 | 0.79 | 1,129 | 0.432 | 0.760 |
| R Parahippocampus | -1.64 | -7.05 | 3.77 | -0.59 | 1,435 | 0.552 | 0.760 |
| L Lateral Orbitofrontal Cortex | -1.23 | -5.22 | 2.75 | -0.61 | 2,111 | 0.544 | 0.760 |
| R Lateral Orbitofrontal Cortex | 0.16 | -5.47 | 5.79 | 0.06 | 598 | 0.955 | 0.993 |
| L Medial Orbitofrontal Cortex | 0.78 | -0.77 | 2.34 | 0.99 | 546 | 0.324 | 0.760 |
| R Medial Orbitofrontal Cortex | 1.07 | -0.60 | 2.75 | 1.25 | 790 | 0.210 | 0.684 |
| L Caudal Middle Frontal Gyrus | 2.28 | -35.06 | 39.63 | 0.12 | 5,245 | 0.905 | 0.980 |
| R Caudal Middle Frontal Gyrus | 30.10 | -9.14 | 69.34 | 1.50 | 5,220 | 0.133 | 0.576 |
| L Rostral Middle Frontal Gyrus | -1.33 | -11.38 | 8.72 | -0.26 | 3,088 | 0.795 | 0.939 |
| R Rostral Middle Frontal Gyrus | 4.70 | -0.23 | 9.64 | 1.87 | 4,616 | 0.062 | 0.555 |
| L Pars Opercularis | -20.47 | -49.72 | 8.77 | -1.37 | 2,373 | 0.170 | 0.632 |
| R Pars Opercularis | 10.97 | -12.53 | 34.48 | 0.91 | 3,891 | 0.360 | 0.760 |
| L Pars Triangularis | -9.66 | -19.89 | 0.56 | -1.85 | 2,920 | 0.064 | 0.555 |
| R Pars Triangularis | 8.48 | -6.60 | 23.55 | 1.10 | 1,463 | 0.271 | 0.704 |
| L Pars Orbitalis | -2.34 | -4.48 | -0.20 | -2.14 | 3,748 | **0.032** | 0.555 |
| R Pars Orbitalis | 0.44 | -1.69 | 2.56 | 0.40 | 3,490 | 0.689 | 0.895 |
| L Superior Frontal Gyrus | 32.42 | -22.63 | 87.47 | 1.15 | 4,433 | 0.248 | 0.704 |
| R Superior Frontal Gyrus | 43.69 | -9.48 | 96.86 | 1.61 | 3,970 | 0.107 | 0.558 |
| L Precuneus | 6.44 | -14.94 | 27.82 | 0.59 | 5,538 | 0.555 | 0.760 |
| R Precuneus | -1.46 | -24.07 | 21.15 | -0.13 | 5,558 | 0.899 | 0.980 |
